# Non-canonical, potassium-driven cerebrospinal fluid clearance

**DOI:** 10.1101/2020.08.03.234260

**Authors:** Huixin Xu, Ryann M Fame, Cameron Sadegh, Jason Sutin, Christopher Naranjo, Della Syau, Jin Cui, Frederick B Shipley, Amanda Vernon, Fan Gao, Yong Zhang, Michael J. Holtzman, Myriam Heiman, Benjamin C Warf, Pei-Yi Lin, Maria K Lehtinen

**Author notes:** These authors contributed equally (Huixin Xu, Ryann M Fame). Bioinformatics Resource Center in the Beckman Institute at Caltech, Pasadena, CA 91125, USA.

## Abstract

Cerebrospinal fluid (CSF) provides vital support for the brain. Abnormal CSF accumulation is deleterious for perinatal neurodevelopment, but how CSF leaves the brain during this critical period is unknown. We found in mice a postnatal neurodevelopmental transition phase featuring precipitous CSF K^+^ clearance, accompanied by water, through the choroid plexus (ChP). The period corresponds to a human fetal stage when canonical CSF clearance pathways have yet to form and congenital hydrocephalus begins to manifest. Unbiased ChP metabolic and ribosomal profiling highlighted this transition phase with increased ATP yield and activated energy-dependent K^+^ transporters, in particular the Na^+^-K^+^-Cl^−^ and water cotransporter NKCC1. ChP-targeted NKCC1 overexpression enhanced K^+^-driven CSF clearance and enabled more permissive cerebral hydrodynamics. Moreover, ventriculomegaly in an obstructive hydrocephalus model was improved by ChP-targeted NKCC1 overexpression. Collectively, we identified K^+^-driven CSF clearance through ChP during a transient but critical neurodevelopmental phase, with translational value for pathologic conditions.

## INTRODUCTION

A balance between cerebrospinal fluid (CSF) production and clearance (influx/efflux) is essential for normal brain function and development (Fame and Lehtinen, 2020). Disrupted CSF volume homeostasis with excessive CSF accumulation is implicated in many pediatric brain disorders, in particular congenital hydrocephalus (Kahle et al., 2016), where patients suffer from a potentially life-threatening accumulation of CSF and frequently develop neurological deficits that last through childhood and into adult life (Vinchon et al., 2012). Schizophrenia patients can have enlarged lateral ventricles by their first episode of psychosis (Steen et al., 2006), and in some cases as early as infancy (Gilmore et al., 2010), suggesting a role for CSF clearance abnormalities in this and possibly other neurodevelopmental disorders. As another example, autism spectrum disorders are associated with altered CSF distribution patterns and enlarged CSF space surrounding the brain (Shen et al., 2017). A better understanding of developing CSF dynamics may help explain why early phases of brain development (e.g. from third trimester to 6 months after birth in human) represent a period of high vulnerability to certain congenial disorders (Volpe, 2008, Gilmore et al., 2010, Shen et al., 2017).

Critically, how CSF is cleared during this perinatal period remains a mystery. Progress in CSF dynamics research has identified several CSF clearance routes including arachnoid granulations, perineural and paravascular pathways, and meningeal lymphatics (Antila et al., 2017, Munk et al., 2019, Fame and Lehtinen, 2020). However, these systems only fully appear at later stages of life (up to 2 years in human and several weeks postnatal in mice) (Antila et al., 2017, Munk et al., 2019), and therefore are not available to contribute to CSF dynamics during these critical early phases. Identifying and manipulating the early endogenous CSF clearance mechanisms could provide one powerful approach for tackling neurodevelopmental disorders involving CSF dysregulation, and may also be applied to fluid disorders affecting adults.

To identify how early CSF is cleared, we investigated tissues with the ability to modulate CSF at this stage. The choroid plexus (ChP) is an intraventricular epithelial structure that forms the majority of the blood-CSF barrier and develops prenatally. It contains diverse ion and fluid transporters along its vast surface area capable of bidirectional transport (Damkier et al., 2013). Although the prevailing model posits that the ChP provides net unidirectional, luminal secretion of ions and water to form CSF, insufficient corroborating data have been collected under physiological experimental conditions. Furthermore, historical clinical observations suggest some absorptive functions of the ChP (Milhorat et al., 1970) which is supported by animal studies (Oreskovic et al., 2017). Finally, broad transcriptional changes of the machinery regulating fluid/ion transport support the concept of temporally dynamic and possibly context-dependent ChP functions in determining net directionality of CSF transport (Liddelow et al., 2013, Delpire and Gagnon, 2019).

To further explore potentially absorptive properties of the ChP, we studied the expression of transporters, the energetic systems and ionic gradients that govern their activity, and their physiological effects across the timespan of early postnatal development in mice. Taken together our data support a novel role and mechanism for CSF clearance by the Na^+^-K^+^-Cl^−^ and water co– transporter, NKCC1, in the apical membrane of the ChP during a specific developmental period. These results have implications for the pathophysiology of congenital disorders accompanied by dysregulated CSF and could inform strategies for treatment of neonatal hydrocephalus and perhaps other disorders.

## RESULTS

### CSF K^+^ declines precipitously during a specific perinatal period

We discovered a unique and transient phase of neurodevelopment when CSF [K^+^] decreased rapidly. We used inductively coupled plasma optical emission spectrometry (ICP-OES) and ion chromatography (IC) to measure levels of key ions likely to govern CSF flux including Na^+^, K^+^, and Cl^−^ at several developmental timepoints. CSF [K^+^] was remarkably high at birth (9.6 ± 3.5 mM), decreased rapidly to 4.4 ± 0.9 mM by P7 (**Fig. 1A**), and later achieved adult levels of 3.1 ± 0.6 mM (**Fig. 1A**) while [Na^+^] and [Cl^−^] were minimally changed (**Fig. 1B**). The reduction in CSF K^+^ was consistent with previous reports in other species (Saunders et al., 2018) and correlated with parallel changes in serum [K^+^] such that the ratio between blood and CSF [K^+^] remained stable (**Fig. 1C**).

**Fig. 1.**
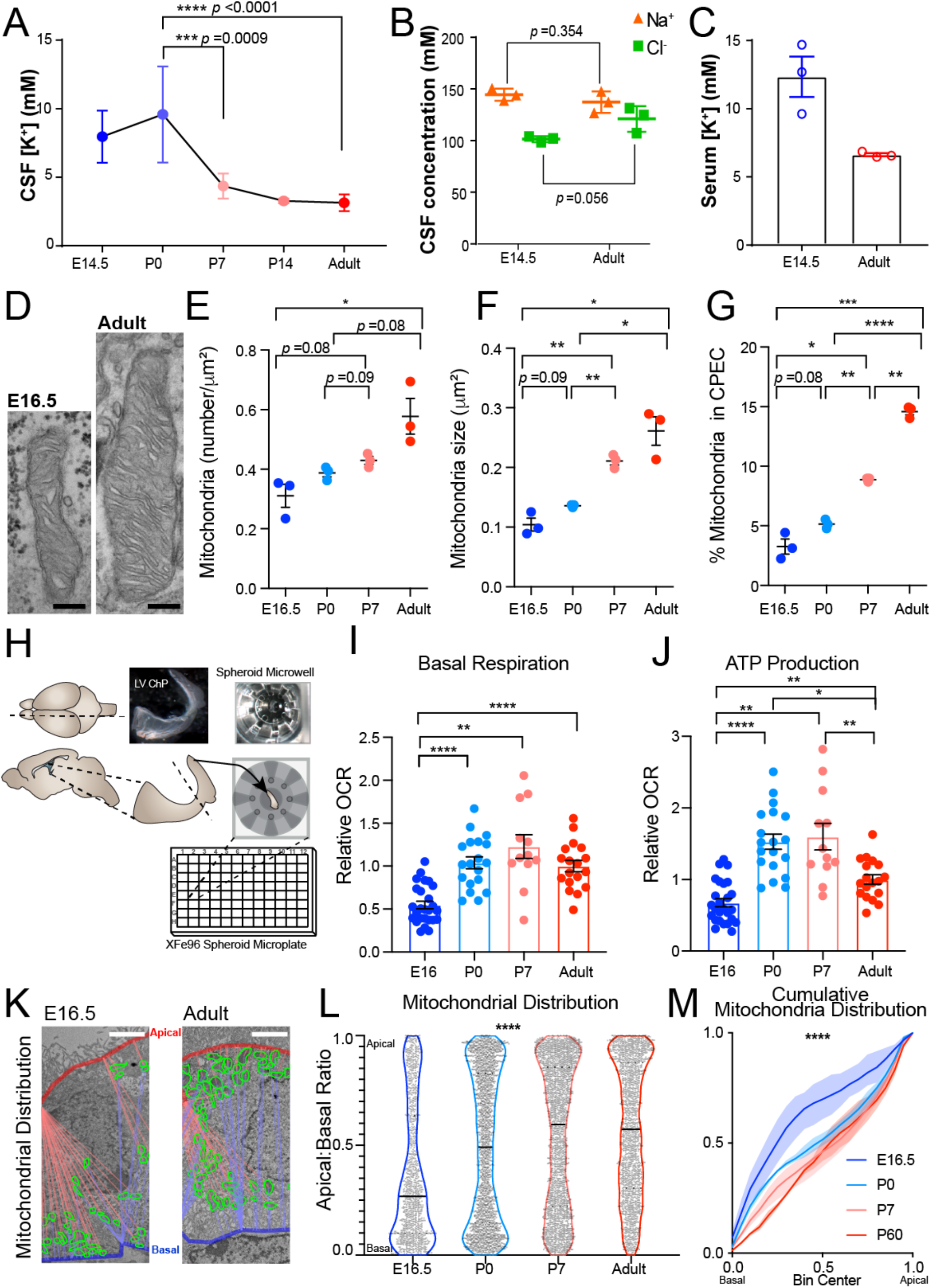
Postnatal CSF [K^+^] decrease coincides with increased ChP metabolism. **(A)** ICP-OES quantification of CSF [K^+^]. N = 6, **** *p* < 0.0001 by one-way ANOVA. Sidak’s test was used for P0 vs. P7 and P0 vs. adult comparison. *** *p* = 0.0009, **** *p* < 0.0001. **(B)** ICP-OES and IC measurements of E14.5 and adult mouse CSF [Na^+^] and [Cl^−^]. N=3; Welch’s t-test. **(C)** ICP-OES measurements of embryo vs. adult mouse serum [K^+^]. N=3. **(D)** Transmission micrographs of mitochondria in LVChP. (**E-G)** Quantification of mitochondrial number (**E**), area (**F**), and % area occupancy (**G**) in ChP epithelial cells. N =3 animals, 5-10 FOV per animal, \**p* < 0.05, \*\**p* < 0.01, \*\*\**p* < 0.005, \*\*\*\**p* < 0.001; Welch’s t-test. **(H)** Schematic of explant-based Agilent Seahorse XFe96 test. (**I-J)** Oxidative respiration metrics over development. \**p* < 0.05, \*\**p* < 0.01, \*\*\*\**p* < 0.0001; Welch’s ANOVA with Dunnett’s T3 multiple comparison test. **(K-L)** Mitochondrial distribution between the apical and basal surfaces (Apical: Basal proximity ratio). 1 = apical surface. 0 = basal surface. Solid line indicates median and dashed line indicates upper/lower quartiles. **** *p* < 0.0001; Kolmogorov–Smirnov test. **(M)** Cumulative distribution of mitochondrial localization. Solid lines are the mean, shaded area represents the range. Scale bar (d) = 250nm, (k) = 2μm. Unless otherwise noted, all quantitative data are presented as Mean ± SEM.

Notably, K^+^ transport has been associated with water co-transport by several K^+^ transporters in various tissues and cell types (Zeuthen, 1994, Hamann et al., 2010, Zeuthen and Macaulay, 2012), suggesting that CSF [K^+^] changes could drive water movement in the brain as well. Therefore, we sought to identify mechanisms underlying this fast clearance of CSF K^+^, which may shed light on CSF outflow during this time.

### ChP metabolism increases during the early postnatal transition phase

We found that the transitional period of rapid CSF K^+^ clearance coincided with high ChP metabolism. We reasoned that K^+^ clearance during this period could be ChP-mediated because the ChP expresses high levels of K^+^ co-transporters on its large CSF-contacting surface area (Keep and Jones, 1990, Damkier et al., 2013). Similar to water and ion transport by other epithelial structures such as kidney proximal and distal tubes (Bhargava and Schnellmann, 2017), K^+^ clearance from CSF by the ChP would be energy-dependent and therefore be accompanied by upregulation of ATP production and mitochondrial activity. Therefore, we evaluated the metabolic status and ATP production capacity of the ChP epithelium before, during, and after the time period of CSF [K^+^] reduction. We found that both mitochondria number and size increased from E16.5 to 2mo (**Fig. 1D-G**), while cellular glycogen load gradually decreased (**Supplementary Fig. 1**). Both observations are consistent with reports from ChP in other mammalian species (Netsky and Shuangshoti, 1975, Keep and Jones, 1990) and suggest functional changes in ChP oxidative metabolism. To assess this we monitored oxygen consumption to calculate basal metabolism and ATP production at embryonic day 16.5 (E16.5), postnatal day 0 (P0), P7, and adult (2 months old (2mo)) ChP explants using Agilent Seahorse XFe technology (**Fig. 1H**, **Supplementary Fig. 2**). We found that E16.5 ChP had the lowest basal respiration of all tested ages (**Fig. 1I**, **Supplementary Fig. 2**). Adult had a higher capacity for overall ATP production than E16.5 ChP, but surprisingly, P0-P7 ChP were the most metabolically active as per ATP production (**Fig. 1I, J**). In addition, mitochondrial subcellular distribution in ChP epithelium was biased toward the apical surface as postnatal development proceeded, with E16.5 mitochondria heavily distributed along the basal side of epithelial cells, P0 mitochondria intermediately localized, and P7 and 2mo mitochondria having more apical distribution (**Fig. 1K-M**, **Supplementary Fig. 3**). Mitochondrial subcellular localization responds to regional energy demand in other cellular processes such as migration of mouse embryonic fibroblasts and during axonal outgrowth (Schuler et al., 2017, Smith and Gallo, 2018). Together with the increase in ATP production postnatally, the shift in ChP epithelial mitochondria distribution over postnatal development suggests increasing ATP supply to meet high demand at the apical ChP surface during the early postnatal phase, concurrent with the rapid clearance of CSF K^+^. This concurrence prompted us to investigate energy-dependent mechanisms whereby ChP epithelial cells might contribute to K^+^ clearance.

### The ChP increases production of CSF-facing ion and water transporters postnatally

Consistent with rapid CSF K^+^ clearance and high ChP metabolism, we found that expression of the energy-dependent cation transport pathway components were upregulated in ChP postnatally. To unbiasedly identify candidates controlling postnatal CSF clearance through the ChP, we conducted ribosomal profiling to investigate transcripts that are prioritized for translation in embryonic (E16.5) and adult (2 mo) ChP, using Translating Ribosomal Affinity Purification (TRAP; Heiman et al., 2008). ChP epithelial cells were targeted by crossing *FoxJ1:cre* mice (Zhang et al., 2007) with *TRAP* (*EGFP:L10a*) mice (Heiman et al., 2008) (**Fig. 2A**, **Supplementary Fig. 4A, B**), and mRNA associated with the L10a ribosomal subunit were purified for sequencing.

**Fig. 2.**
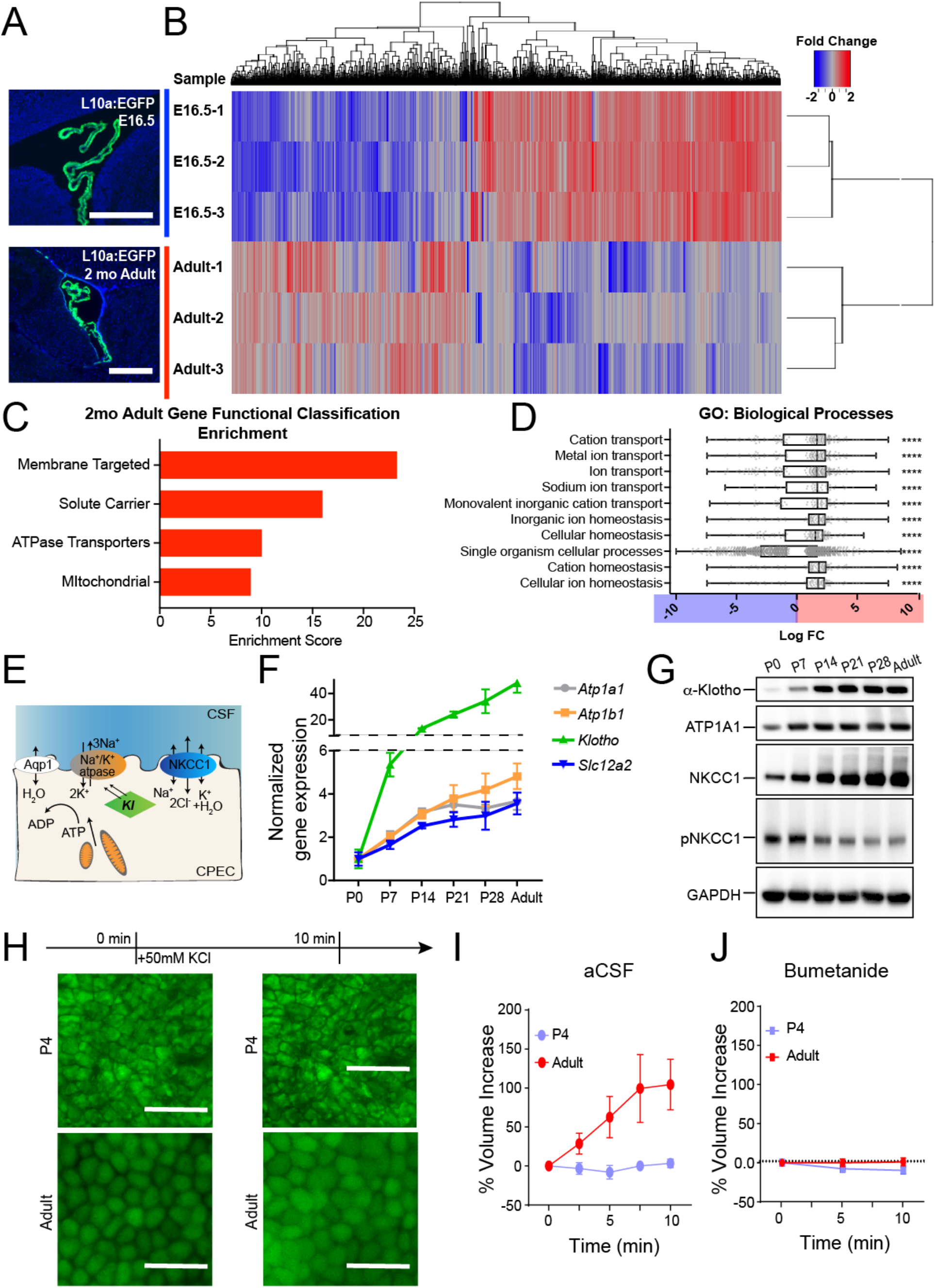
Choroid plexus epithelial cells display age-dependent translation of ion and water transporters, in particular NKCC1. **(A)** Rpl10a-conjugated EGFP expression in ChP epithelial cells after *Foxj1*-Cre recombination in TRAP-BAC mice. **(B)** Heatmap and hierarchical clustering of differentially expressed genes (adjusted p < 0.05). **(C)** Top 4 gene functional clusters shown by DAVID to be enriched in Adult ChP epithelial cells over E16.5 ChP epithelial cells. **(D)** Top 10 significantly enriched gene ontology (GO) terms for “Biological processes”. Plotted with boxes for quartiles and whiskers for 5^th^ and 95^th^ percentiles. The log_10_ fold change (LogFC) is plotted for each expressed gene for the network. Positive values (red) indicate Adult enrichment and negative values (blue) indicate E16.5 enrichment. *p* values are corrected for multiple measures using Bonferroni correction. \*\**p* ≤0.01, \*\*\**p* ≤0.001, \*\*\*\**p* ≤0.0001. **(E)** Schematics demonstrating the interaction of NKCC1, Na^+^/K^+^-ATPase, and Klotho on the apical membrane of a ChP epithelial cell (CPEC). **(F-G)** RT-qPCR and immunoblotting of LVChP during postnatal development. **(H)** Fluorescence images of Calcein-AM labeled epithelial cells from LVChP explants under high extracellular K^+^ challenge. Scale bar = 50μm. **(I-J)** Quantification of ChP epithelia cellular volume by IMARIS 3D analysis. Percent volume increase = dV/V_0_ for each timepoint (t). V_0_ = initial volume of the cell; t = subsequent timepoint after addition of challenge; dV= V_t_ − V_0_ % 100%. At least 10 cells were analyzed for each explant from one animal; N = 4.

TRAP analyses revealed 1967 differentially translated transcripts (adjusted *p* < 0.05) between E16.5 and 2mo adult ChP: 1119 enriched at E16.5 and 847 enriched at 2mo (**Fig. 2B**). Gene set and pathway analyses revealed developmentally regulated ChP programs. Adult (2mo) ChP had enriched functional gene sets associated with active transmembrane membrane transport and mitochondria, which is consistent with our abovementioned findings on metabolism changes (**Fig. 2C**). Specifically, cation transport was enriched, supporting the hypothesis of ChP mediating CSF K^+^ transport postnatally (**Fig. 2C, D**). Enriched pathways in the 2mo adult included secretion associated pathways named for other, better studied secretory processes, including salivary and pancreatic secretion, all of which have a special emphasis on water and ion transmembrane transport (**Supplementary Fig. 4C (red), D**). Consistent with a rise in fluid and ion modulating machinery, there was a striking enrichment of more transmembrane and signal peptide-containing transcripts in adult ChP (**Supplementary Fig. 4E, F**). These results indicate that the ChP specifically gained fluid and ion modulatory functions postnatally.

### NKCC1 is poised to mediate perinatal ChP CSF K^+^ and water clearance

Among all fluid and ion modulating candidates with increasing postnatal expression (**Supplementary Fig. 4G, H**), we identified NKCC1 (*Slc12a2*) as the candidate most likely to mediate CSF clearance. NKCC1 is functionally related to Na^+^/K^+^-ATPase (*Atp1a1* and *Atp1b1*), as the latter actively maintains the Na^+^/K^+^ gradient that powers NKCC1. Both Na^+^/K^+^-ATPase and NKCC1 are capable of CSF K^+^ clearance, but NKCC1 was of particular interest because (1) it is a co-transporter of K^+^ and water (Zeuthen and Macaulay, 2012, Steffensen et al., 2018); and (2) the activity of NKCC1 can be further modified by phosphorylation (Darman and Forbush, 2002), lending additional control to its fluid/ion modulatory capacity. In addition, NKCC1 is particularly enriched in the ChP and does not impact broad functionality like the Na^+^/K^+^-ATPase does, both of which are ideal features for a functional therapeutic intervention target. We refined our temporal expression analyses of NKCC1, ATP1a1, ATP1b1, and Klotho (*Kl*), which contributes to the membrane localization of the Na^+^/K^+^-ATPase (Razzaque, 2008, Sopjani et al., 2011) (**Fig. 2E**), by sampling weekly from P0 to P28 and then at 2mo and confirmed increased expression of transcript and protein for each component across developmental time (**Fig. 2F, G**, **Supplementary Fig. 5**). The observed changes in NKCC1 total protein were corroborated by an independent approach where the rate of ChP epithelial cell swelling under high [K^+^] challenge (Steffensen et al., 2018) reflected the abundance of NKCC1 protein (**Fig. 2H-J**, **Supplementary Fig. 6**).

In addition, we found particularly high levels of phosphorylated, therefore activated (Darman and Forbush, 2002), NKCC1 (pNKCC1) in the ChP of P0-P7 pups, with P7 having peak pNKCC1 levels among all postnatal ages, indicative of increased NKCC1 activity during the first postnatal week (**Fig. 2G**). Similar to the timeline of ChP ATP production (**Fig. 1J**), the timeframe of high ChP pNKCC1 was concurrent with the fast CSF [K^+^] decrease during the first postnatal week (**Fig. 1A**), suggesting a functional correlation and further confirming the significance of the early postnatal transitional period. Taken together, we identified ChP NKCC1 as the top candidate for mediating postnatal CSF K^+^ and water clearance.

### NKCC1 temporal regulation requires epigenetic control that is implicated in congenital hydrocephalus

We found that the temporal profile of NKCC1 expression was tightly regulated at the epigenetic level by modulators implicated in some forms of congenital hydrocephalus. The NuRD complex governs differentiation and maturation of diverse cells and tissues (Goodman and Bonni, 2019). Our previously published RNA sequencing studies (Lun et al., 2015) identified NuRD components, including the ATPase CHD family members (Chd4 being the most highly expressed), the histone deacetylases HDAC1/2, and methyl CpG-binding domain protein MBD3 in the ChP (**Fig. 3A**). *De novo* loss-of-function CHD4 mutations are implicated in some groups of children with congenital hydrocephalus and ventriculomegaly (Weiss et al., 2020). We found that CHD4 localized to nuclei in mouse ChP epithelial cells beginning at P0 (**Fig. 3B**). Immunoprecipitation of CHD4 identified HDAC1, HDAC2, and MBD3 by immunoblotting in mouse ChP (**Fig. 3C**, technical control for Co-IP protocol is shown in **Supplementary Fig. 7**), confirming the existence of the CHD4/NuRD complex in developing ChP. We then disrupted the complex by generating ChP-*Chd4* deficient mice. Cre was expressed in *Chd4* floxed mice (*Chd4* fl/fl) (Williams et al., 2004) using an adeno-associated viral vector (AAV) with tropism for the ChP (AAV2/5) (Haddad et al., 2013), delivered by *in utero* intracerebroventricular (ICV) injection at E14.5. *Chd4* transcript levels dropped to <50% by P7 (**Fig. 3D**). While CHD4 protein levels only substantially decreased by P14 (**Fig. 3E, G**), we found that the developmental increase of ChP NKCC1 expression was disrupted as soon as the CHD4 protein decreased and lasted at least until P28 (**Fig. 3F, G**). Similar results were also observed in 4VChP (**Fig. 3H, I**). The expression of other developmentally regulated, functionally relevant candidates (*atp1a1, atp1b1, and klotho*) was not affected (data not shown). These data confirm that the NuRD/ChD4 complex is one of the required components tightly regulating ChP NKCC1 developmental expression.

**Fig. 3.**
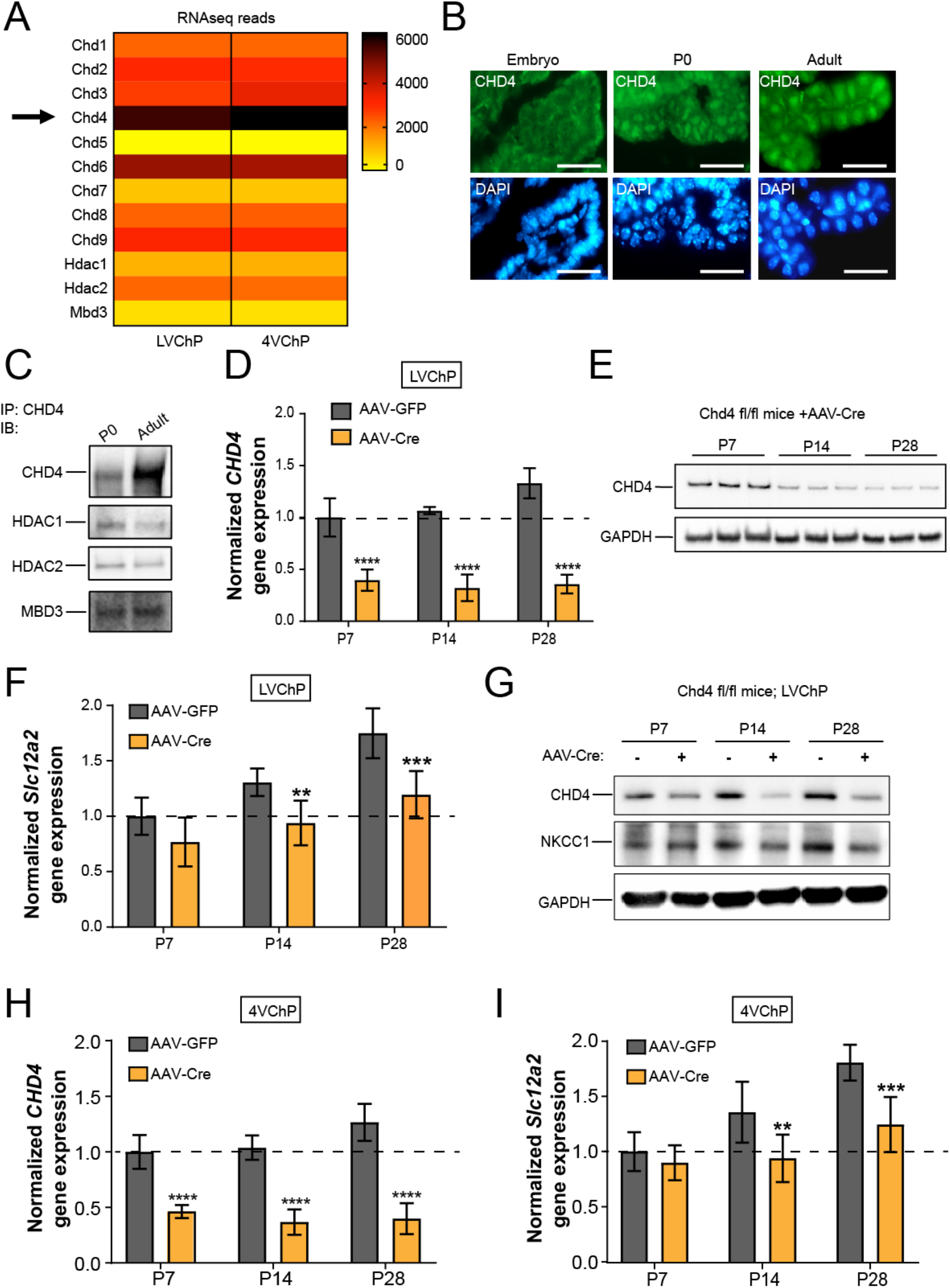
NKCC1 temporal expression requires CHD4/NuRD complex. **(A)** RNAseq data showing expression of CHD and other NuRD units by the ChP. **(B)** Immunofluorescence images of CHD4 in the ChP epithelia at E16.5, P0, and adult; Scale bar = 30μm. **(C)** Immunoblots of Co-IP by CHD4 antibody. **(D)** RT-qPCR of CHD4 transcripts in ChP with AAV2/5-Cre infection.**** *p*< 0.0001, N = 7, Welch’s t-test. **(E)** Immunoblot of CHD4 in AAV-cre mice ChP lysate with. **(F)** RT-qPCR of NKCC1 expression in AAV-cre vs. AAV-GFP mice ChP. All values were normalized to P7 AAV-GFP control mice. \*\**p* = 0.0015, \*\*\**p* < 0.001, N = 7, Welch’s t-test. **(G)** Immunoblot of NKCC1 in LVChP lysates from AAV-cre vs. AAV-GFP mice. **(H-I)** CHD4 and NKCC1 RT-qPCR in 4VChP. ** p = 0.0083, *** *p* < 0.001, **** *p* < 0.0001, N = 7, Welch’s t-test.

### ChP NKCC1 actively mediates CSF clearance during the early postnatal transition phase

To test whether NKCC1 is indeed capable of transporting from CSF into the ChP during the period of rapid CSF [K^+^] decline, we induced NKCC1 overexpression (OE) in developing ChP epithelial cells using AAV2/5. NKCC1 transport directionality follows combined Na^+^, K^+^, Cl^−^ gradients, which are close to being neutral in adult brains and likely to bias towards the CSF-to-ChP direction during the early postnatal phase. NKCC1 protein level would be rate-limiting during the early postnatal time when it is already highly phosphorylated, unlike in older mice where pNKCC1 only represented a small portion of total NKCC1. The goal of this OE approach was to accelerate endogenous ChP NKCC1 transport, thereby revealing its directionality based on whether CSF [K^+^] clearance was enhanced or delayed. AAV2/5-NKCC1, which expresses NKCC1 fused to an HA tag (Somasekharan et al., 2013), or control GFP virus was delivered by *in utero* ICV at E14.5. Successful NKCC1 OE and increased pNKCC1 was confirmed in ChP at P0 (**Fig. 4A-D**). Appropriate localization to apical membranes of epithelial cells, transduction efficiency, and tissue specificity were also validated (**Fig. 4E-I**, **Supplementary Fig. 8 A, B**). Transcript levels of other K^+^ transporters or channels did not change following AAV2/5-NKCC1 transduction (**Supplementary Fig. 8C**). Because CSF [K^+^] sharply decreased from P0 to P7 (**Fig. 1A**), we sampled CSF from ChP NKCC1 OE and control mice at P1. We found that ChP NKCC1 OE reduced CSF [K^+^] more than controls, with their P1 CSF [K^+^] values closely approximating those normally observed at P7 (**Fig. 4J**), indicating accelerated K^+^ clearance from CSF after enhanced ChP NKCC1 activity. CSF total protein levels were not affected (AAV2/5-GFP = 2.50 ± 0.20 mg/ml vs. AAV2/5-NKCC1 = 2.71 ± 0.46 mg/ml; N = 6 from two litters each; *p* = 0.34, unpaired t-test). Overall, these findings support a model in which, under physiological conditions with high early postnatal CSF [K^+^], ChP NKCC1 transports K^+^ out of CSF.

**Fig. 4.**
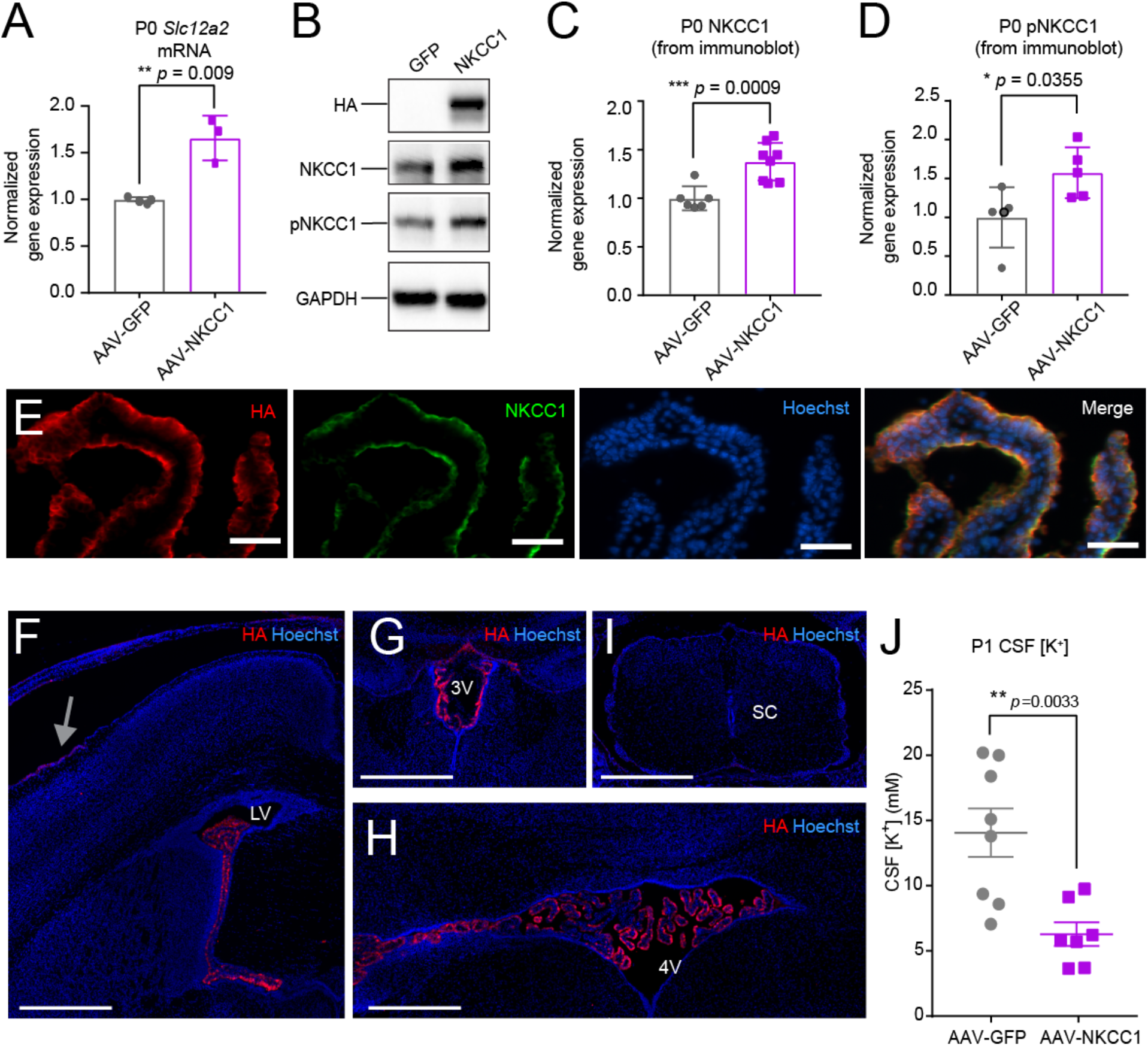
ChP NKCC1 actively mediates CSF K^+^ clearance in the first postnatal week. **(A)** RT-qPCR of NKCC1 mRNA levels in P0 mice. ** *p* = 0.009, N=3; Welch’s t-test. **(B)** Immunoblots from AAV-NKCC1 vs. AAV-GFP P0 mice ChP lysates. **(C-D)** Quantification of all immunoblots of NKCC1 (**C**) *** *p* = 0.0009, N=7; Welch’s t-test; and pNKCC1 (**D**) * *p* = 0.0355, N=5; Welch’s t-test. **(E)** Immunofluorescence images showing co-localization of 3xHA tag and NKCC1 in P0 ChP. Scale bar = 50μm. **(F-I)** Immunofluorescence images of HA in AAV2/5-NKCC1 transduced brain at P1: the LVChP (**F**), 3^rd^ ventricle ChP (3VChP; **G**), 4VChP (**H**), and the spinal cord (**I**; sc = spinal cord). Traces of HA is shown in the meninges near the injection site (grey arrow). Scale bar = 500μm. **(J)** ICP-OES measurements of CSF [K^+^] from AAV-NKCC1 vs. AAV-GFP P1 mice (N = 8 in AAV-GFP cohort; N = 7 in AAV-NKCC1 cohort). \*\**p* = 0.0033; Welch’ t-test.

Next, we found that the circulating CSF volume in ChP NKCC1 OE mice was reduced, as reflected by smaller lateral ventricles. To avoid any tissue processing artifacts, we conducted live T2-weighted magnetic resonance imaging (MRI) (**Fig. 5A**) to quantify lateral ventricle volume. AAV-GFP mice were indistinguishable from naive wild-type mice at P14. In contrast, NKCC1 OE mice had reduced lateral ventricle volumes (**Fig. 5A, B**), without decrease in overall brain size (**Fig. 5C**), reflecting less circulating CSF. The difference in ventricle sizes from these same mice was sustained up to our final measurement at P50 (AAV-GFP: 3.12 ± 0.59 mm^3^ vs. AAV-NKCC1: 1.28 ± 0.28 mm^3^, * *p* = 0.0182). While the exact transport direction of NKCC1 in adult ChP is still under debate (Delpire and Gagnon, 2019), the consistency in ventricular volume from P14 into later life supports our working model that because a relatively small proportion of ChP NKCC1 was phosphorylated in mice P14 and older (**Fig. 2G**), NKCC1 levels are not rate-limiting and thus OE would not as substantially impact ChP functions in older animals. Collectively, our findings demonstrate that ChP NKCC1 mediated CSF clearance during the first postnatal week. Augmenting this process impacted CSF volume homeostasis in the long term.

**Fig. 5.**
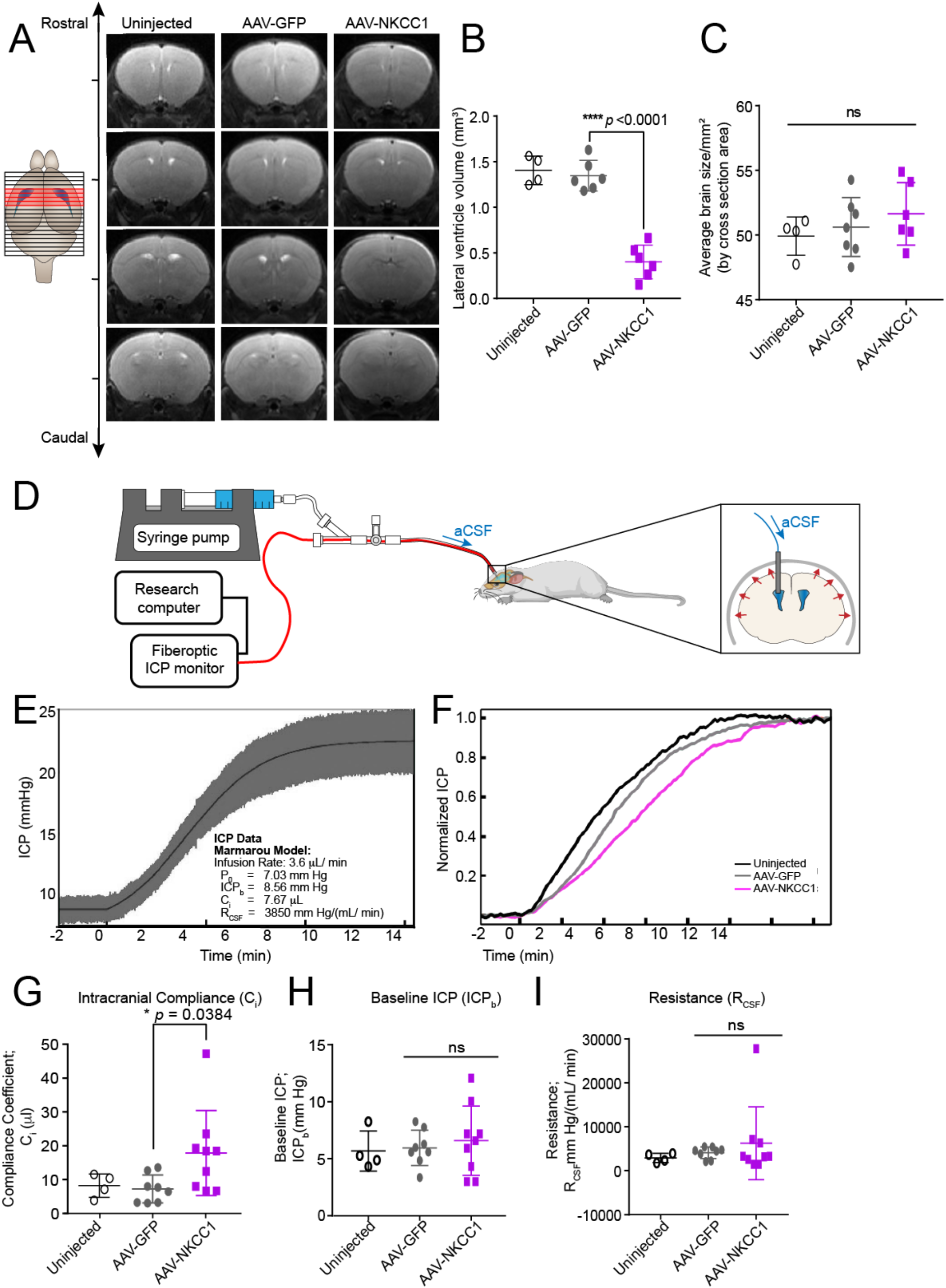
ChP NKCC1 overexpression reduces brain ventricular volume and increases intracranial compliance. **(A)** T2-weighted live MRI images. Only slices with visible LV are shown (marked red in the schematics). **(B)** LV volumes. Uninjected N = 4; AAV-GFP and AAV-NKCC1 N = 6; \*\*\*\**p* < 0.0001; Welch’s t-test. **(C)** Brain sizes, which are presented as the average coronal section area from all images with visible ventricles (NKCC1 OE data were calculated using the matching images to the controls, regardless of ventricles visibility). Uninjected N = 4; AAV-GFP and AAV-NKCC1 N = 6; Welch’s t-test. **(D)** Schematic of *in vivo* constant rate CSF infusion test. **(E)** Example of ICP curve during the infusion test (infusion begins at 0 min) in an AAV-GFP mouse, fitted to Marmarou’s model. Values extracted include: baseline ICP (*ICP_b_*), a pressure-independent compliance coefficient (*C_i_*) and the resistance to CSF outflow (*R_CSF_*). **(F)** Example ICP recordings from AAV-NKCC1 mice and controls. For clarity, data have been low pass filtered to remove the waveform components. **(G)** Compliance coefficients. Uninjected N = 4; AAV-GFP N = 8; AAV-NKCC1 N = 9; \**p* = 0.0384; Welch’s t-test. **(H-I)** Plots of baseline ICP and resistance to CSF outflow (R_CSF_) at 5-7 weeks. Uninjected N=4, AAV-GFP N=8, AAV-NKCC1 N=9; Welch’s t-test.

We then tested and found that enhancing CSF clearance through ChP NKCC1 OE changed how the brain and cranial space adapted to CSF volume changes. Intracranial compliance (C_i_) and CSF resistance (R_CSF_) describe the ability of the entire intracranial space (including brain, meninges, and outflow routes) to accommodate an increasing CSF volume that would otherwise increase intracranial pressure (ICP). In humans, these parameters are measured by a CSF constant rate infusion test (Aquilina et al., 2012, Eide, 2018, Lalou et al., 2018) and can aid in diagnosis and evaluation of conditions like hydrocephalus, which has decreased C_i_ (Kahle et al., 2016). We developed a miniaturized version of this test to determine the C_i_ and R_CSF_ in mice. The constant rate infusion test artificially increases CSF volume by ICV infusion of artificial CSF (aCSF), causing ICP to rise and plateau at a new level (**Fig. 5D, E**). The C_i_ and R_CSF_ are estimated from the ICP vs. time curve using Marmarou’s model of CSF dynamics (Czosnyka et al., 2012) (**Supplementary Fig. 9A**). Simply put, the C_i_ is proportional to the rate of ICP increase, and the R_CSF_ is related to the level of the post-infusion ICP plateau (**Fig. 5E**). As a quality control for the correct placement of infusion and measurement catheter, arterial and respiratory pulsations were clearly visible in the ICP waveform and their amplitude increased with volume load as expected (**Supplementary Fig. 9B, C**). Using this approach, we found that ChP NKCC1 OE significantly increased C_i_ at an age of 5-7 weeks (**Fig. 5F, G**), consistent with the brain having greater capacity for CSF in ventricles “deflated” due to excessive CSF clearance. Resting ICP and R_CSF_ were unchanged (**Fig. 5H, I**).

### Enhanced ChP NKCC1 function mitigates ventriculomegaly in a model of obstructive hydrocephalus

Our findings of enhanced CSF clearance after ChP NKCC1 OE indicate that ChP NKCC1 can remove excess CSF. Therefore, we hypothesized that ChP NKCC1 OE expression could mitigate ventriculomegaly in a model of postnatal obstructive hydrocephalus. We first overexpressed ChP NKCC1 at E14.5 by in utero AAV2/5 ICV, then introduced obstructive hydrocephalus by a single unilateral injecting of kaolin into the lateral ventricle at P4 (Shaolin et al., 2015), and finally evaluated the lateral ventricle volumes by live T2 MRI at P14 (**Fig. 6A**). While both NKCC1 OE and control mice had enlarged ventricles at P14, NKCC1 OE mice had reduced ventriculomegaly compared to controls, with the average ventricle volume being less than 1/3 of the controls (**Fig. 6B-D;** ventricles marked by blue arrows; kaolin deposits marked by red arrows). Taken together, our findings demonstrate that early, ChP targeted NKCC1 OE has a sustained and broad impact on specific volumetric and biophysical parameters of the intracranial space with potential therapeutic applications to hydrocephalus.

**Fig. 6.**
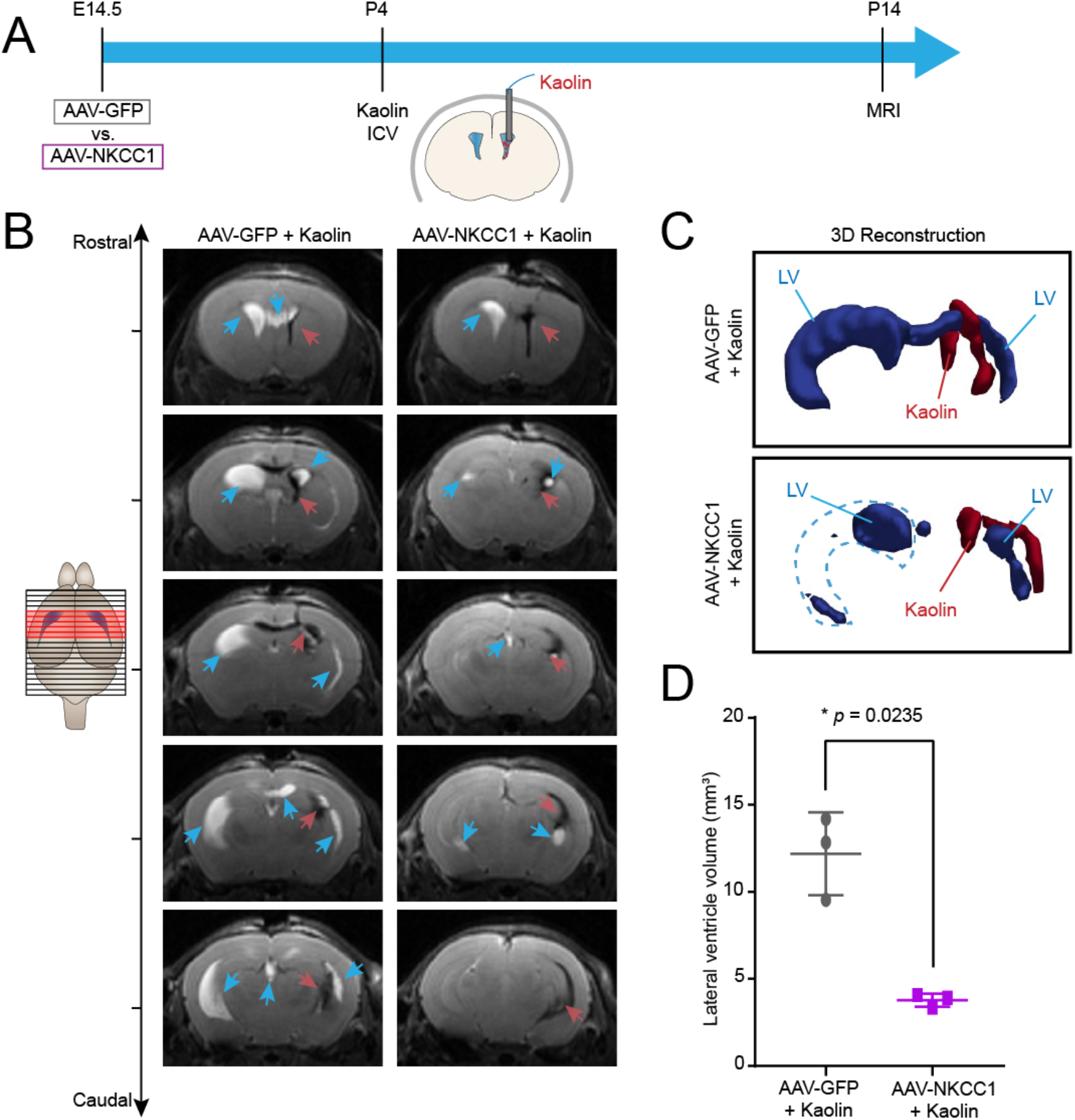
ChP NKCC1 overexpression mitigates ventriculomegaly in an obstructive hydrocephalus model. **(A)** Schematics showing the workflow: E14.5 in utero ICV of AAV2/5-NKCC1 or AAV2/5-GFP, followed by ICV of kaolin at P4, and MRI at P14. **(B)** Representative sequential brain images (rostral to caudal) by T2-weighted live MRI images. Blue arrows: LV. Red arrows: kaolin. **(C)** 3D reconstruction of the LV and kaolin deposition. LV: blue. Kaolin: red. **(D)** LV volumes. N = 3; \**p* = 0.0.0235; Welch’s t-test.

## DISCUSSION

In this study we sought to understand how CSF is cleared from the brain before the development of canonical CSF outflow routes (e.g. arachnoid granulations and meningeal lymphatics) (Antila et al., 2017, Munk et al., 2019). The intervening time period is a critical, transient phase in brain development when failure of CSF clearance has debilitating consequences (Volpe, 2008). Our results suggest that this period is defined by rapid decrease in CSF K^+^. The ChP mediates the CSF K^+^ clearance during this transition period, and thus forms a CSF outflow route through ion and water co-transport by NKCC1 (**Fig. 7**). This CSF clearance by the ChP contrasts the prevailing models that ChP constantly, unidirectionally secretes CSF. Taken together, we discovered an unconventional, precisely timed, function of the developing ChP that clears CSF prior to the formation of other canonical routes, and provides targets for fluid management intervention during a critical transition phase of brain development.

**Fig. 7.**
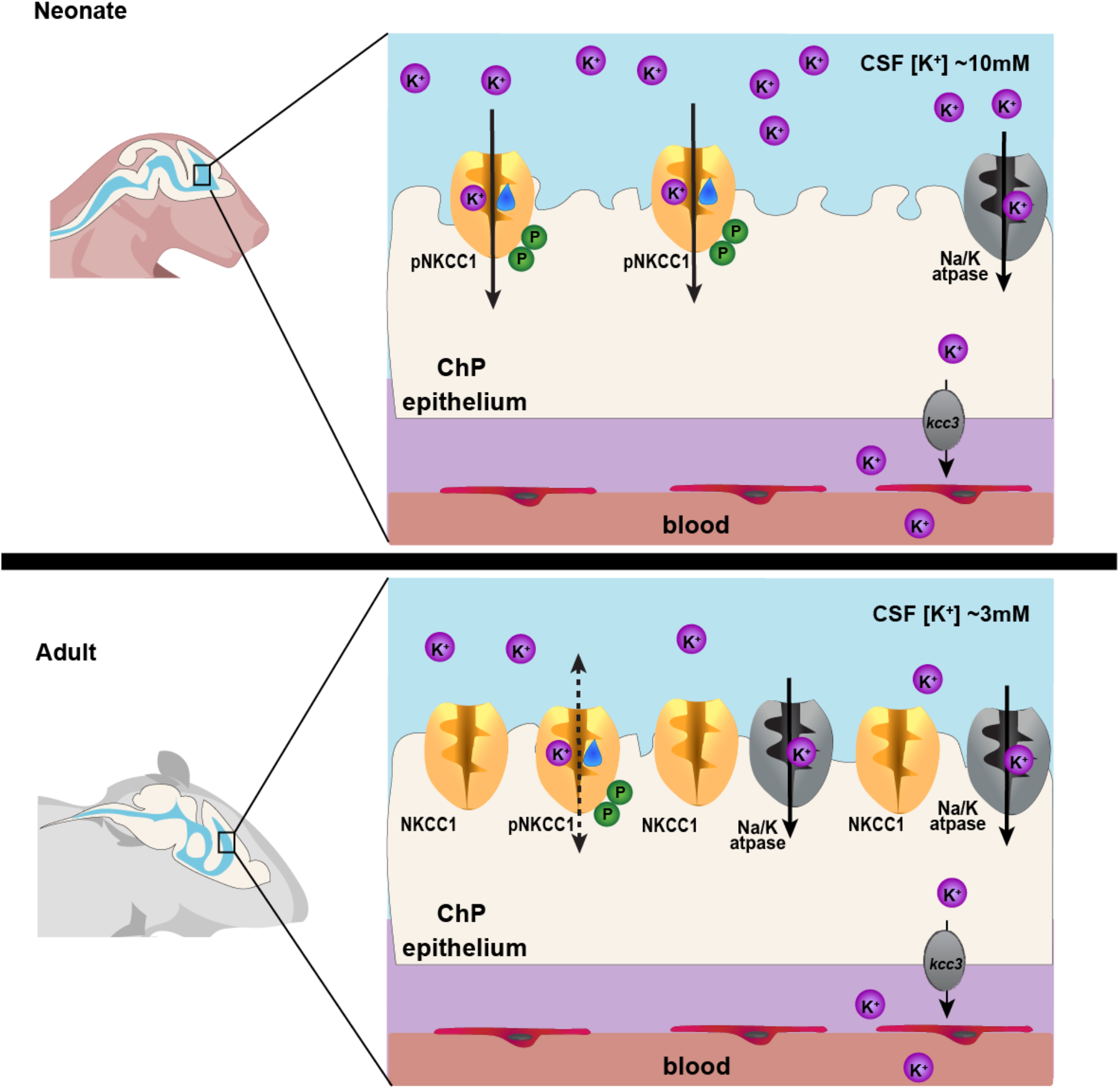
Working model of ChP NKCC1 mediating K^+^-driven CSF outflow. The schematics describes ChP NKCC1 mediated K^+^ and water clearance from CSF in neonatal mice, in comparison to the adult scenario. For simplicity and clarity, only K^+^ is depicted among all ions and only NKCC1 and Na^+^/K^+^-ATPase are included. Neonatal (P0-7, left) ChP has high pNKCC1 than adult, albeit lower total NKCC1. Neonate CSF [K^+^] is 2-3 fold higher than adult. With similar [Na^+^] and [Cl^−^], this [K^+^] difference is sufficient to alter the total Nernst potential of epithelial cells and bias NKCC1 transport of K^+^, together with water, out of CSF into the ChP in neonates.

NKCC1 is a bidirectional transporter, recently discovered to be an important co-transporter of water in the adult ChP (Steffensen et al., 2018). Although clearly established as a key molecular mechanism of CSF regulation, ChP NKCC1 transport direction and its determinants *in vivo* have been actively debated due to the technical challenges of 1) specifically manipulating ChP NKCC1 without affecting NKCC1 in other CSF-contacting cells, instead of ICV application of chemicals such as NKCC1 inhibitor bumetanide; and 2) accurately determining intracellular ion levels of ChP epithelial cells, and therefore ion gradients, under physiological conditions, as summarized in **Supplementary Table 1** and reviewed by Delpire and Gagnon (Delpire and Gagnon, 2019). Our *in vivo* “gain-of-function” approach effectively bypasses the abovementioned technical limitations. By overexpressing NKCC1 specifically in the ChP through AAV transduction to amplify its physiological functional impact, we could subsequently observe the resulting CSF K^+^ and fluid volume changes to reveal the transporter’s native directionality. Using this approach, we found that, in contrast to the common notion that the ChP constantly produces CSF, NKCC1 in the ChP mediated CSF clearance when CSF [K^+^] is above adult values, especially during the first postnatal week in mice. This phase corresponds to the third trimester to 6 months after birth in human, which represents a window of high vulnerability to congenial fluid disorders (Volpe, 2008).

We next demonstrated that the ChP clearance of CSF can be targeted to temper abnormal CSF accumulation. The ChP has been targeted for therapeutic manipulation in rodent models of neurologic diseases ranging from Huntington’s disease and lysosomal storage disorders, to Alzheimer’s disease, where transduction of exogenous gene products into ependymal or ChP epithelial cells has improved cardinal symptoms of disease (Kaler, 1994, Hudry et al., 2013). Encouragingly, we showed that enhancing ChP epithelial cell NKCC1 transport capacity lessened the severity of ventriculomegaly in a model of obstructive hydrocephalus. Our data demonstrate the possibility of tempering congenital hydrocephalus by augmenting endogenous ChP NKCC1 activity to increase CSF absorption rates during early development when CSF [K^+^] is high. In addition, because ChP CSF absorption via NKCC1 is driven by increased CSF [K^+^], this mechanism may come into play in other pathogenic conditions where CSF [K^+^] is transiently increased, such as after tissue injury or ventricular bleeding. Therefore, our findings emphasize the ChP as a targetable, K^+^-sensitive and on-demand CSF drainage route in neurological disorders where CSF homeostasis is disrupted.

Further, in light of recent findings reporting hydrocephalus and ventriculomegaly in children with *de novo* loss-of-function CHD4 mutations (Weiss et al., 2020), we found that the CHD4/NuRD complex is required for the developmental regulation of NKCC1 expression. This connection suggests a possible pathophysiological mechanism whereby lack of CHD4 activity might reduce NKCC1 levels during early development (equivalent to P0-P7 in mice), and lead to insufficient CSF clearance resulting in hydrocephalus. In our loxP-cre approach, most CHD4 protein knockdown and resulting stagnation of NKCC1 expression occurred by P14, which is beyond the critical window of NKCC1 activity at P0-P7. As such, we did not model developmental ventriculomegaly with this approach. Improved genetic tools for early CHD4 knockout and new animal models harboring the *de novo* patient mutations would be required to fully unravel the regulatory connection between CHD4/NuRD complex and NKCC1.

A key question that emerges from this work is: If the ChP is acting as an outflow pathway rather than a source of CSF during this transitional developmental phase – where does the early CSF water content come from? One mechanism that could be acting at this stage is CSF secretion by the developing brain tissue (e.g. progenitor cells that have a cell body at the ventricular zone but extend their basal processes to the developing pia) which secretes CSF immediately after neural tube closure (Gato et al., 2014). Future studies should elucidate whether this mechanism extends into this transitional phase. Consistent with progenitor involvement in CSF dynamics, recent identification of genes driving pediatric hydrocephalus shows affected genes to be expressed predominantly by cortical progenitor cells lining the brain’s ventricles (Furey et al., 2018), and not the ChP, suggesting a non-choroidal source of CSF as an alternative contributor to abnormal CSF production.

In addition to fluid regulation, the newly identified ChP clearance route provides a key mechanism to regulate extracellular K^+^ during the first postnatal week. The subunits of the major system for moving K^+^ against its individual concentration gradient, the Na^+^/K^+^-ATPase (i.e. Atp1a1 and Atp1b1), were not yet at their full expression levels during this period. The ChP NKCC1-mediated K^+^ clearance mechanism might assist in establishing the K^+^ gradient in a timely manner, which is crucial for cellular physiology (Rasmussen et al., 2020). Notably, the period of rapidly decreasing CSF [K^+^] overlaps with the developmental phase when the excitatory-to-inhibitory “GABA switch” occurs. In early cortical progenitor cells that reside in the ventricular zone and are bathed by CSF, the classic inhibitory neurotransmitter GABA leads to excitatory potentials and suppression of DNA synthesis (LoTurco et al., 1995). As newborn cortical neurons differentiate and migrate away from the ventricular zone, GABA switches to adopt the more classic role as an inhibitory neurotransmitter by lowering intracellular Cl^−^ (Owens et al., 1999) which is achieved through coordinated activities of neuronal K^+^/Cl^−^ co-transporters KCC2 and NKCC1 (Pisella et al., 2019, Watanabe et al., 2019). Because ions, including K^+^, can traffic from CSF into interstitial fluid (Cserr, 1965, Fencl et al., 1966), any interference with the developmental timeline of ChP NKCC1 that resulted in delayed CSF K^+^ clearance could potentially increase extracellular/interstitial fluid [K^+^] and affect neural physiology (Rasmussen et al., 2020). Specifically, such a change in extracellular/interstitial fluid [K^+^] could fundamentally impact neuronal NKCC1 and KCC2 transport equilibrium, potentially contributing to a delayed GABA switch, a phenomenon reported in many models of neurodevelopmental and psychiatric disorders including subtypes of autism spectrum disorder (Amin et al., 2017), Rett syndrome (Banerjee et al., 2016), Fragile X syndrome (He et al., 2014), schizophrenia (Hyde et al., 2011), and Down syndrome (Deidda et al., 2015). Furthermore, extracellular [K^+^] and certain K^+^ channels also regulate activities of microglia (Madry et al., 2018), which are critical in synaptic pruning during postnatal neurodevelopment in mice (Schafer et al., 2012). Thus, the ChP is poised to play important roles in proper CNS formation by creating and maintaining desirable extracellular ionic homeostasis at different developmental stages, with subsequent effects on neuronal maturation, circuit formation, and neuroinflammatory homeostasis.

Beyond key findings and implications during this critical transitional developmental stage, our study introduced a murine ICP measurement device combined with constant CSF infusion. This approach provides a much-needed advance in fluid research technology that can be broadly applied to study essentially all CSF dynamic systems across the mouse lifespan. We adapted our tool from clinical practice to provide a range of options for measuring global cerebral fluid states that reflect the interaction between CSF and cranial tissues. In later life, CSF homeostasis is maintained by collaborative efforts from multiple players in the brain, including the ChP, the dural lymphatics (Antila et al., 2017), glymphatics (Munk et al., 2019), leptomeningeal vasculature (Li et al., 2020), and the ependyma (Spassky et al., 2005). While this approach measures overall cranial fluid dynamics as one single unit, future applications could apply mathematical models that have been proposed to isolate the contribution of distinct CSF outflow routes using data acquired from human patients (Vinje et al., 2020). Such adaptability secures the broadening relevance of our tool and inspires optimism for further improved resolution in studying brain fluid dynamics. Availability of this new tool also allows future researchers to obtain measurements in support of the growing comprehensive “systems” view of regulatory mechanisms of CSF-brain interactions.

In summary, our study presents a critical transient phase when the ChP acts as a non-canonical route for CSF clearance prior to the maturation of other canonical clearance pathways. ChP NKCC1 mediates CSF clearance in a K^+^-dependent manner. Targeting this absorption route holds promise in improving fluid management for congenital hydrocephalus and other CSF disorders.

## ACKNOWLEDGEMENTS

We thank members of the Lehtinen, Heiman, and Warf labs for helpful discussions; Nancy Chamberlin for critical reading of the manuscript; Katia Georgopoulos for sharing the *Chd4* fl/fl mouse line and associated genotyping methods; P. Ellen Grant for the ICP monitor. We thank the following facility and personnel: Maria Ericsson and HMS EM facility; Yaotang Wu and Michael Marcotrigiano and BCH Small Animal Imaging Laboratory; the MIT BioMicro Center (TRAP sequencing); BCH viral core and University of Pennsylvania Vector Core.

## Author contributions

H.X., R.M.F., C.S. J.S., P.-Y. L., B.C.W., F.B.S., J.C., D.S., C.N., and M.K.L. designed and performed experiments; H.X., R.M.F., C.S., and J.S. analyzed the data; Y.Z. and M.J.H. provided material; A.V., F.G., and M.H. provided technological support; H.X., R.M.F., and M.K.L. wrote the manuscript. All co-authors edited the manuscript.

## METHODS

### Mice

The Boston Children’s Hospital IACUC approved all experiments involving mice in this study. Timed pregnant CD1 dams were obtained from Charles River Laboratories. Mice with germline loxP-CHD4-loxP were imported from MGH and bred in-house. Both male and female mice were equally included in the study and were analyzed at postnatal day 0, 7, 14, 21, 28, 5-7weeks, and 2+ months. Animals were housed in a temperature-controlled room on a 12-hr light/12-hr dark cycle and had free access to food and water. For studies involving mice younger than postnatal day 10, all mice were allocated into groups based solely on the gestational age without respect to sex (both males and females were included). For studies involving mice older than 10 days, both male and female are included intentionally.

### CSF Collection and Metal Detection

CSF was collected by from cisterna magna using a glass capillary, and collected CSF was centrifuged at 10,000xg for 10min at 4℃ to remove any tissue debris. Metal quantifications were performed by Galbraith Laboratories, Inc. (Knoxville, TN, USA). Inductively coupled plasma - optical emission spectrometry (ICP-OES) was used for K and Na quantification, and ion chromatography (IC) was used for the Cl^−^ quantification. All tests were performed using 5-7μL of CSF.

### TRAP

Mice aged 8 weeks or E16.5 from the *Foxj1*:Cre x EGFP-L10a Bacterial Artificial Chromosome (BAC) transgenic lines (N=3, each N included LVChP pooled from 3 mice) were used and brain tissue was immediately dissected and used for TRAP RNA purifications as previously described (Heiman et al., 2008). RNA quality was assessed using Bioanalyzer Pico Chips (Agilent, 5067-1513) and quantified using Quant-iT RiboGreen RNA assay kit (Thermo Fisher Scientific R11490). Libraries were prepared using Clonetech SMARTer Pico with ribodepletion and Illumina HiSeq to 50NT single end reads. Sequencing was performed at the MIT BioMicroCenter.

### Sequencing Data Analysis

The raw fastq data of 50-bp single-end sequencing reads were aligned to the mouse mm10 reference genome using STAR 2.4.0 RNA-Seq aligner (Dobin et al., 2013). The mapped reads were processed by htseq-count of HTSeq software (Anders et al., 2015) with mm10 gene annotation to count the number of reads mapped to each gene. The Cuffquant module of the Cufflinks software (Trapnell et al., 2010) was used to calculate gene FPKM (Fragments Per Kilobase of transcript per Million mapped reads) values. Gene differential expression test between different animal groups was performed using DESeq2 package (Love et al., 2014) with the assumption of negative binomial distribution for RNA-Seq data. Genes with adjusted p-value< 0.05 are chosen as differentially expressed genes. All analyses were performed using genes with FPKM> 1, which we considered as the threshold of expression (Figure 2-source data).

### Sequencing Pathway and Motif Analysis

Functional annotation clustering was performed using DAVID v6.7 (Huang da et al., 2009). Gene ontology (GO) analysis was performed using AdvaitaBio iPathway guide V.v1702. Enrichment vs. perturbation analysis was performed by AdvaitaBio iPathway guide V.v1702 and allows comparison of pathway output perturbation and cumulative gene set expression changes. In brief, the enrichment analysis is a straightforward gene-set enrichment over representation analysis (ORA) considering the number of differentially expressed genes (DEGs) that are assigned to a given pathway. The enrichment value is expressed as a proportion of enriched members to total genes in a defined pathway and a p-value (Fisher) is calculated for this score, however false positives have been reported at up to 10% with this method (Draghici et al., 2007). Perturbation, on the other hand, uses pathway data that applies relationships between gene products rather than only using a list. Perturbation assigns an impact score based on a mathematical model that captures the entire topology of the pathway and uses it to calculate how changes in the expression of each gene in the pathway would perturb the absolute output of the pathway (Draghici et al., 2007). Then, these gene perturbations are combined into a total perturbation for the entire pathway and a p-value is calculated by comparing the observed value with what is expected by chance. Motif analyses were performed using SignalP (v5.0; Almagro Armenteros et al., 2019) and TMHMM (v2.0; Sonnhammer et al., 1998).

### Transmission Electron Microscopy

All tissue processing, sectioning, and imaging was carried out at the Conventional Electron Microscopy Facility at Harvard Medical School. Forebrain tissues were fixed in 2.5% Glutaraldehyde/2% Paraformaldehyde in 0.1 M sodium cacodylate buffer (pH 7.4). They were then washed in 0.1M cacodylate buffer and postfixed with 1% Osmiumtetroxide (OsO4)/1.5% Potassiumferrocyanide (KFeCN_6_) for one hour, washed in water three times and incubated in 1% aqueous uranyl acetate for one hour. This was followed by two washes in water and subsequent dehydration in grades of alcohol (10 minutes each; 50%, 70%, 90%, 2×10min 100%). Samples were then incubated in propyleneoxide for one hour and infiltrated overnight in a 1:1 mixture of propyleneoxide and TAAB Epon (Marivac Canada Inc. St. Laurent, Canada). The following day, the samples were embedded in TAAB Epon and polymerized at 60 degrees C for 48 hours. Ultrathin sections (about 80nm) were cut on a Reichert Ultracut-S microtome, and picked up onto copper grids stained with lead citrate. Sections were examined in a JEOL 1200EX Transmission electron microscope or a TecnaiG² Spirit BioTWIN. Images were recorded with an AMT 2k CCD camera.

### Glycogen and Mitochondrial Quantification

Glycogen and mitochondrial quantification was performed by hand using the ImageJ plugin FIJI (Schindelin et al., 2012, Schneider et al., 2012). Percentages were calculated by dividing the area of interest by the total area of ChP epithelial cell within the field of view. No other cell types were included in the analysis. For each condition, analyses were performed across multiple individual animals (N=3 for each age). From each animal, 10-20 fields of view were imaged at 3,000x for glycogen analysis and 5-10 fields of view were imaged at 3,000x for mitochondrial analysis. Each different field of view represented a unique cell or cells, and fields of view were chosen such that both the apical and basal surfaces of the cell were visible. For mitochondrial distribution, a custom MatLab (v.2018) code was written to extract the centroid from mitochondria data traced in ImageJ ROIs (Supplementary file). Then a distance transformation was performed from each mitochondrion centroid to the hand-traced apical or basal surfaces. The shortest distance was extracted to calculate the apical: basal proximity ratio, such that 1= on the apical surface and 0= on the basal surface. The analyses included a total of 1747 adult mitochondria, 2241 P7 mitochondria, 2257 P0 mitochondria, and 1123 embryonic mitochondria.

### Seahorse Metabolic Analysis

ChP explants were dissected in HBSS (Fisher, SH30031FS) and maintained on wet ice until plated. Only the posterior leaflet of the P0, P7, and adult ChP was retained for analysis due to empirically determined limitations of the oxygen availability in the XFe96 Agilent Seahorse system. Tissue explants were plated on Seahorse XFe96 spheroid microplates (Agilent, 102905-100) coated with Cell TAK (Corning), in Seahorse XF Base Medium (Agilent, 102353-100) supplemented with 0.18% glucose, 1mM L-glutamine, and 1mM pyruvate at pH7.4 and incubated for 1 hour at 37 °C in a non-CO_2_ incubator. Extracellular acidification rates (ECAR) and oxygen consumption rates (OCR) were measured via the Cell Mito Stress Test (Agilent, 103015-100) with a Seahorse XFe96Analyzer (Agilent) following the manufacturer’s protocols. Data were processed using Wave software (Agilent). ATP production was calculated as the difference in OCR measurements before and after oligomycin injection, as described by the manufacturer’s protocol (Agilent, 103015-100). The Cell Mito Stress test was performed 2-5 independent times. The individual analyses were performed by averaging the readings from both the right and left hemisphere lateral ventricle ChP for each individual. Data were normalized by Calcein-AM (2μM in PBS, Life Technologies L-3224) fluorescence measured at the end of the assay. Data are presented normalized to the adult levels for each assay to account for any experimental variability.

### High K^+^ challenge study

Fresh LV ChPs were collected from P4 pups and adult mice in room temperature HBSS and glued down onto imaging dishes with coverslip bottom. The tissues were incubated at 37 °C with Calcein-AM (Invitrogen L3224; 1:200) for 10min and then rinsed with 37 °C artificial CSF (aCSF: 119 mM NaCl, 2.5 mM KCl, 26 mM NaHCO_3_, 1 mM NaH_2_PO_4_, 11 mM glucose, with fresh 2.0 mM magnesium chloride and 2.8 mM calcium chloride). The tissues were soaked in 1.8ml aCSF at the beginning of each imaging session and allowed to stabilize for 10min. One Z-stack was acquired to reflect the baseline cell volume. Then a 10x KCl solution in aCSF was spiked into the bath to make the final bath K^+^ concentration 50mM immediately before imaging subsequent continued. A total of five 3D Z-stacks were acquired throughout a 10-min imaging session to capture changes in cellular volume over time. Each stack took less than 30s to minimize changes in cell volume from the beginning to the end of each stack. All imaging studies were carried out at 37°C. Image stacks were imported into Imaris (Bitplane) software. Individual epithelial cells were identified by shape. Cells with discrete borders that were present at all timepoints and had dark pixels both above and below them in Z for the whole timecourse were selected *a priori* and then traced by hand using the “Surpass” functionality to create a 3D surface volume through all Z stacks based on Calcein-AM uptake signal. Due to known z-step distance and interpolation between the planes, Imaris calculated the number of voxels for each cell. This analysis was then repeated for the same cell throughout the timecourse. We verified manually that the cell was the same individual based on the topology of the surrounding cells, allowing for adjustment for any x-y drifting that occurred. The relative volume was calculated as dV/V_0_ for each timepoint (t) where V_0_ is the initial volume of the cell, t is each subsequent timepoint after addition of challenge, and dV= V_t_-V_0_.

### Tissue processing

Samples were fixed in 4% paraformaldehyde (PFA). For cryosectioning, samples were incubated in the following series of solutions: 10% sucrose, 20% sucrose, 30% sucrose, 1:1 mixture of 30% sucrose and OCT (overnight), and OCT (1 hour). Samples were frozen in OCT.

### Immunostaining

Cryosections were blocked and permeabilized (0.3% Triton-X-100 in PBS; 5% serum), incubated in primary antibodies overnight and secondary antibodies for 2 hours. Sections were counterstained with Hoechst 33342 (Invitrogen H3570, 1:10000) and mounted using Fluoromount-G (SouthernBiotech). The following primary antibodies were used: chicken anti-GFP (Abcam ab13970; 1:1000), mouse anti-Aqp1 (Santa Cruz sc-32737; 1:100), rabbit anti-CHD4 (Abcam ab72418, 1:200), rabbit anti-NKCC1 (Abcam ab59791; 1:500), rat anti-HA (Roche 11867423001; 1:1000). Secondary antibodies were selected from the Alexa series (Invitrogen, 1:500). Images were acquired using Zeiss LSM880 confocal microscope with 20x objective.

### Co-IP

Tissues were homogenized in NET buffer (150mM NaCl, 10mM Tris 8.0, 5mM EDTA, 10% glycerol and 2% Triton-100) supplemented with protease inhibitors. Protein concentration was determined by BCA assay (Thermo Scientific 23227). Lysates with same amount of total protein (250-1000μg based on experiments) were pre-cleared at 4° C for 2hr with Protein G agarose and then incubated with desired antibody or control antibody at 4° C overnight (no beads present during antibody incubation). Protein G agarose beads were added to lysate-antibody mixture after overnight incubation for 2hr. Beads were washed thoroughly and then eluted by boiling in 2% SDS. ChP tissues were pooled across 7 litters of P0 pups and 30 adults to achieve sufficient protein for Co-IP.

### Immunoblotting

Tissues were homogenized in RIPA buffer supplemented with protease and phosphatase inhibitors. Protein concentration was determined by BCA assay (Thermo Scientific 23227). Samples were denatured in 2% SDS with 2-mercaptoethanol by heating at 37°C (for NKCC1) or 95°C (for CHD4 and other NuRD complex proteins) for 5 minutes. Equal amounts of proteins were loaded and separated by electrophoresis in a 4-15% gradient polyacrylamide gel (BioRad #1653320) or NuPAGE 4-12% Bis-Tris gel (Invitrogen #NP0322), transferred to a nitrocellulose membrane (250mA, 1.5 hours, on ice), blocked in filtered 5% BSA or milk in TBST, incubated with primary antibodies overnight at 4°C followed by HRP conjugated secondary antibodies (1:5000) for 1 hour, and visualized with ECL substrate. For phosphorylated protein analysis, the phospho-proteins were probed first, and then blots were stripped (Thermo Scientific 21059) and reprobed for total proteins. For co-IP protein analysis, TrueBlot secondary antibody (eBioscience 18-8816-33) was used to detect only non-denatured IgG and avoid background signal from IP antibody. The following primary antibodies were used: rabbit anti-NKCC1 (Abcam ab59791; 1:1000), rabbit anti-pNKCC1 (EMD Millipore ABS1004; 1:1000), rabbit anti-ATP1a1 (Upstate C464.6/05-369; 1:250, goat-anti-klotho (R&D AF1819-sp; 1:200), rabbit anti-GAPDH (Sigma G9545; 1:10000), mouse anti-HA (Abcam ab130275; 1:1000), rabbit anti-CHD4 (Abcam ab72418; 1:2000), rabbit anti-MBD3 (Abcam ab157464; 1:1000), rabbit anti-HDAC1 (Abcam ab7028; 1:2000), mouse anti-HDAC2 (Abcam 51832; 1:2000).

### Quantitative RT-PCR

For mRNA expression analyses, the ChP were collected and pooled from 2 pups. RNA was isolated using the MirVana miRNA isolation kit (Invitrogen AM1561) following manufacturer’s specifications without miRNA enrichment step. Extracted RNA was quantified spectrophotometrically and 100ng was reverse-transcribed into cDNA using the High Capacity cDNA Reverse Transcription kit (Applied Biosystems #4368814) following manufacturer’s specifications. RT-qPCRs were performed in duplicate using Taqman Gene Expression Assays and Taqman Gene Expression Master Mix (Applied Biosystems) with GAPDH as an internal control. Cycling was executed using the StepOnePlus Real-Time PCR System (Invitrogen) and analysis of relative gene expression was performed using the 2^−ΔΔCT^ method. Technical replicates were averaged for their cycling thresholds and further calculations were performed with those means.

### *In utero* intracerebroventricular injection (ICV)

Timed pregnant mice (E14.5) were deeply anesthetized by isoflurane and placed on warm pads. Laparotomy was performed and AAV solution was delivered into the lateral ventricle of each embryo using glass capillary pipettes. The abdominal incision was then sutured. Meloxicam analgesia was longitudinally delivered according to IACUC protocol.

### Intraventricular kaolin injection in postnatal pups

Postnatal day 4 pups (P4) were deeply anesthetized by hypothermia. 1μl of sterile kaolin solution (25% in PBS) was delivered into the left lateral ventricle using glass capillary pipettes. The lateral ventricle location was determined as in between bregma and lambda, and 1mm from mid-line. The pups were then warmed and returned to the dam.

### AAV production

The original AAV-NKCC1 plasmid was purchased from Addgene (pcDNA3.1 HA CFP hNKCC1 WT (NT15-H) was a gift from Biff Forbush: Addgene plasmid # 49077; http://n2t.net/addgene:49077; RRID:Addgene_49077). The plasmid carries an 3xHA tag at the N-terminal of NKCC1 to allow detection and separation from endogenous NKCC1. The CFP tag was removed by BsaI digestion to reduce insert size for AAV production. Virus production and purification were performed by the Penn Vector Core. Due to the very large size of the plasmid we experienced variable infection efficiency. All mice receiving AAV-NKCC1 were analyzed for HA expression after every experiment to confirm infection efficiency. AAV-GFP and AAV-Cre were purchased from BCH viral core at Boston Children’s Hospital.

### Magnetic resonance imaging (MRI)

Mice were imaged using Bruker BioSpec small animal MRI (7T) at 2wk and P50 while under anesthesia by isoflurane. A warm pad was used to maintain body temperature. Breathing rate and heart rate were monitored to reflect the depth of anesthesia. All axial T2 images were acquired using the following criteria: TE/TR=60/4000; Ave=8; RARE=4; slice thickness=0.6mm. Ventricle volumes were calculated by manual segmentation using FIJI/ImageJ. In studies with unilateral kaolin injection, 3D reconstruction of the ventricles was performed by manual segmentation in ITK-SNAP (Madan, 2015) and exported through ParaView.

### Constant rate CSF infusion test (ICP and compliance measurement)

An apparatus was developed to perform a constant infusion test in mice through a single catheter for both infusion of CSF and monitoring of ICP. A 20cc syringe was filled with aCSF and placed in an automated infusion pump (GenieTouch, Kent Scientific Co., Denver) and set to deliver a constant rate infusion of 1-4 uL/minute. The syringe was connected via pressure tubing to hemostasis valve Y connector (Qosina, NY). A fiberoptic ICP sensor (Fiso Technologies Inc, Québec, Canada) was inserted through the other port of the rotating hemostat and then into 0.55 mm diameter catheter until the sensor was flush with the catheter’s distal tip. The entire apparatus and tubing was carefully screened to ensure the absence of air bubbles. Adult mice were then deeply anesthetized, placed on a warm pad, and head-fixed with ear bars. The distal end of the infusion device (catheter with fiberoptic sensor) was placed inside lateral ventricle (− 0.4mm (anterior-posterior) and 1.2mm (medial-lateral) with respect to Bregma, and a depth of 2 mm from the outer edge of the skull); the catheter was then sealed with Vetbond (3M, Minnesota). Intraventricular access and water-tight seal was confirmed by observation of arterial and respiratory waveforms in the ICP signal and a transient rise in ICP upon gentle compression of the abdomen and thorax. Two minutes of baseline ICP were recorded before initiating the infusion of aCSF. As the infusion proceeded, careful observation was made of the mouse’s respiratory rate. After the ICP level reached a new plateau, the infusion was discontinued. The procedure was terminal. Parameters of the Marmarou model of CSF dynamics for constant rate infusions were estimated by a non-linear least squares fit of the model to the ICP data (Czosnyka et al., 2012)

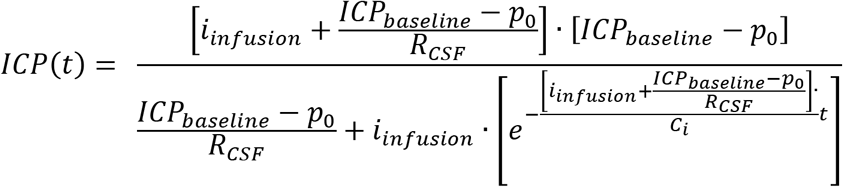

where i_infusion_ is the rate of infusion, *ICP_baseline_* is the ICP level before infusion, *p_0_* is a pressure in the storage arm of the model, *R_CSF_* is the resistance to CSF outflow, and *C_i_* is the compliance coefficient.

## QUANTIFICATION AND STATISTICAL ANALYSIS

Biological replicates (N) were defined as samples from distinct individual animals, analyzed either in the same experiment or within multiple experiments, with the exception when individual animal could not provide sufficient sample (i.e. CSF), in which case multiple animals were pooled into one biological replicate and the details are stated in the corresponding figure legends. Statistical analyses were performed using Prism 7 or R. Outliers were excluded using ROUT method (Q = 1%). Appropriate statistical tests were selected based on the distribution of data, homogeneity of variances, and sample size. The majority of the analyses were done using One-way ANOVA with multiple comparison correction (Sidak) or Welch’s unpaired t-test, except for Fig. 1E-G, and Fig. 1I-J where the analysis was done by Welch’s ANOVA with Dunnett’s T3 multiple comparison test, and Fig. 1L where the analysis was done using Kolmogorov-Smirnov test. F tests or Bartlett’s tests were used to assess homogeneity of variances between data sets. Parametric tests (t-test, ANOVA) were used only if data were normally distributed and variances were approximately equal. Otherwise, nonparametric alternatives were chosen. Data are presented as means ± standard deviation (SD). If multiple measurements were taken from a single individual, data are presented as means ± standard errors of the mean (SEMs). Please refer to figure legends for sample size. *p* values < 0.05 were considered significant (* *p* < 0.05, ** *p* < 0.01, *** *p* < 0.001, **** *p* < 0.0001). Exact p values can be found in the figure legends. P values are also marked in the figures where space allows.

