## Supplementary figures and tables for "Non-canonical, potassium-driven cerebrospinal fluid clearance"

[‡]These authors contributed equally.

*Supplementary Figure 1*

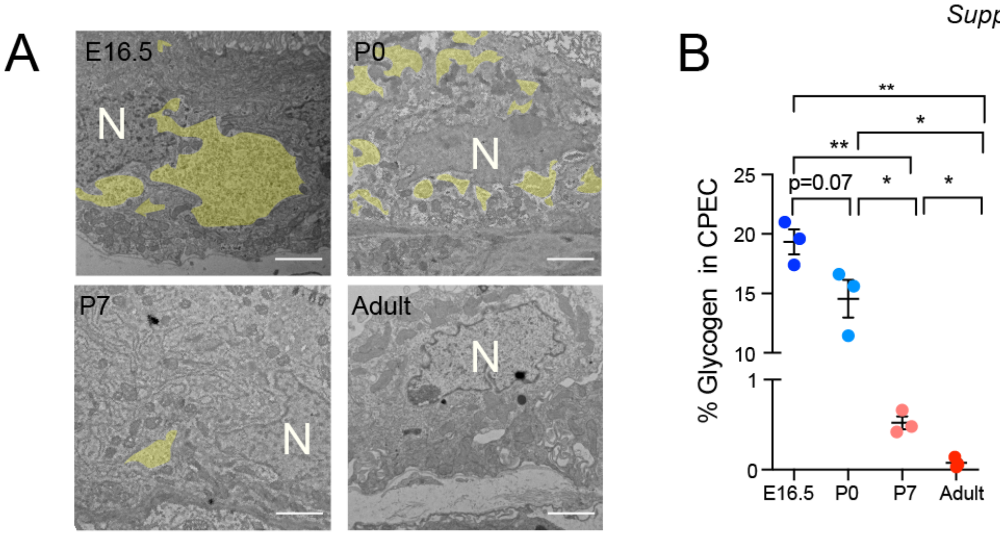


**Supplementary Fig 1. Glycogen load in the ChP epithelial cells. (A)** Representative transmission electron micrographs of E16.5, P0, P7, and adult LV ChP. Glycogen granules are highlighted in yellow. Scale bar = 2μm. N = Nucleus. (**B)** Proportion of TEM fields of view that are filled with glycogen granules, N=3 animals, 10-15 FOV per individual, *$p < 0.05$, **$p < 0.01$; Welch's t-test.

*Supplementary figure 2*

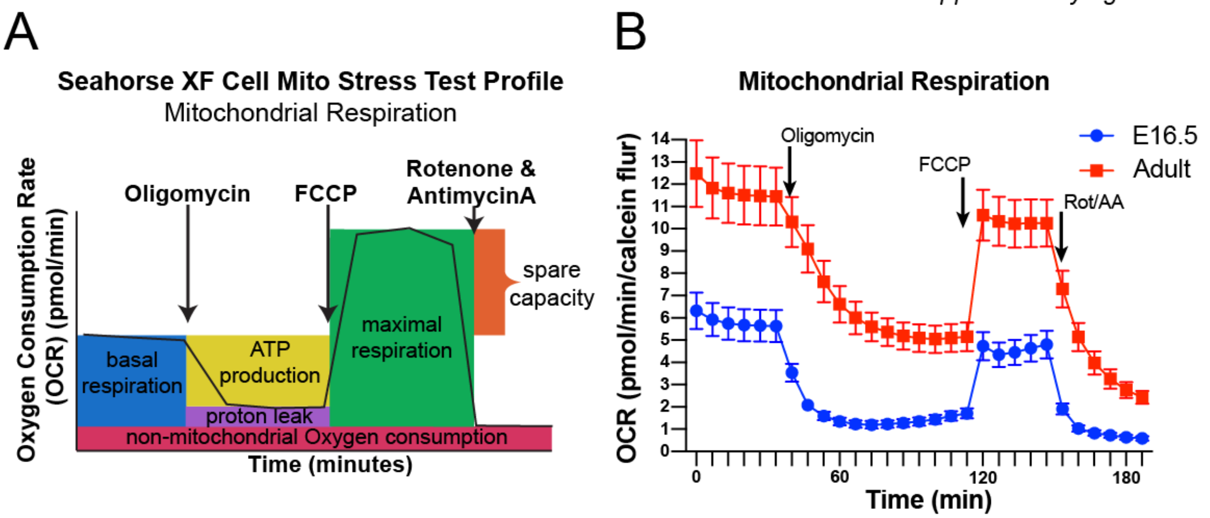


**Supplementary Fig 2. Seahorse XF Cell mito stress test profile and representative curves.** **(A)** Schematic of the Agilent Cell Mito Stress Test showing the experimental design to quantify mitochondria basal respiration and ATP production. (**B**) Representative experiment of ChP in Cell Mito Stress Test; N = 12 E16.5 animals and N = 4 adult animals.

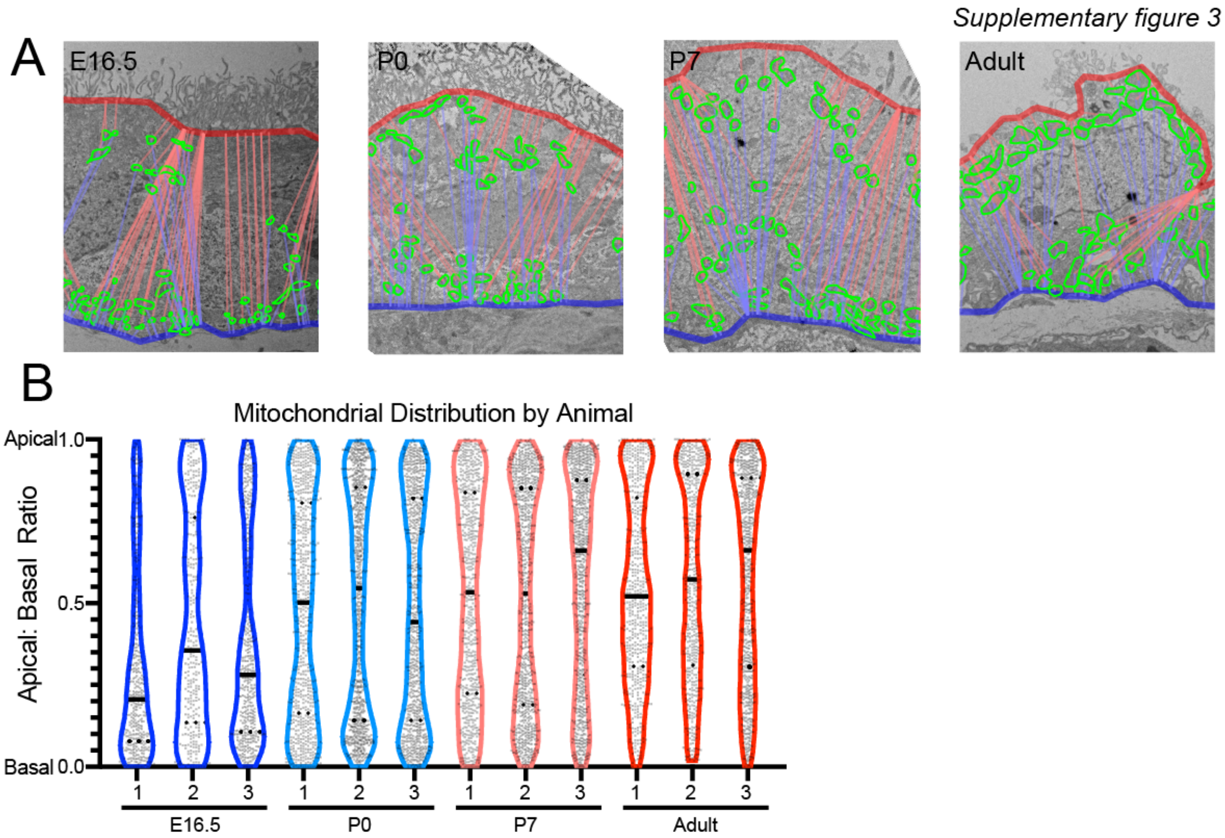


**Supplementary Fig 3. Representative images and quantification of ChP epithelial mitochondria distribution analysis. (A)** Representative transmission electron micrographs of E16.5, P0, P7, and adult LV ChP. Mitochondria (green circle), apical membrane (red line), and basal membrane (blue line) are labeled. **(B)** Mitochondrial distribution plots from each animal. Apical: Basal ratio: 1 is touching the apical surface and 0 is touching the basal surface. Solid line indicates median and dashed line indicates upper/lower quartiles.

*Supplementary figure 4*

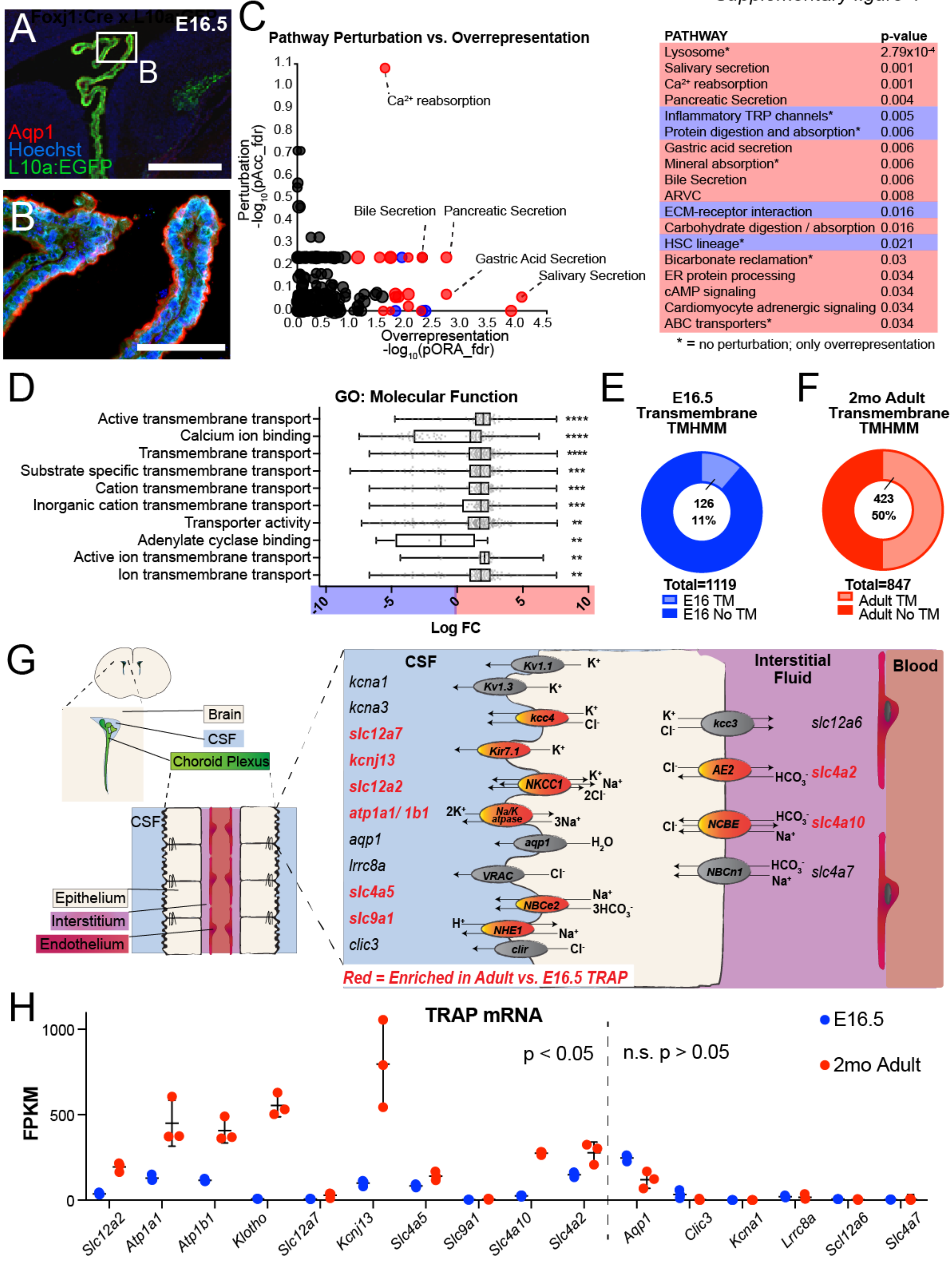

**Supplementary Fig 4. Supportive analysis of TRAP sequencing. (A-B)** Rpl10a-conjugated EGFP expression in ChP epithelial cells after *Foxj1*-Cre recombination in TRAP-BAC mice. Aqp1 marked ChP epithelial apical membrane. **(C)** Perturbation vs. overrepresentation analysis via iPathway (Advaita) reveals enriched pathways at E16.5 (blue) and Adult (red). * indicates pathways that are only overrepresented, but not predicted to be additionally perturbed at the network level. **(D)** Top 10 significantly enriched GO terms for "molecular function". Plotted with boxes for quartiles and whiskers for $5^{th}$ and $95^{th}$ percentiles. The $\log_{10}$ fold change (LogFC) is plotted for each expressed gene for the network. Positive values (red) indicate Adult enrichment and negative values (blue) indicate E16.5 enrichment. *p* values are corrected for multiple measures using Bonferroni correction. $^{**}p \leq 0.01$, $^{***}p \leq 0.001$, $^{****}p \leq 0.0001$. **(E-F)** Proportion of enriched genes in E16.5 (blue) and Adult (red) ChP with predicted transmembrane domains using TMHMM. **(G)** Schematic of ChP localization within brain ventricles, relative position to blood and CSF, and transporters. Red highlights: significantly enriched in Adult vs. E16.5 TRAP (adjusted $p < 0.05$). **(H)** FPKM values from TRAP of transcripts associated with ChP transport.

*Supplementary figure 5*

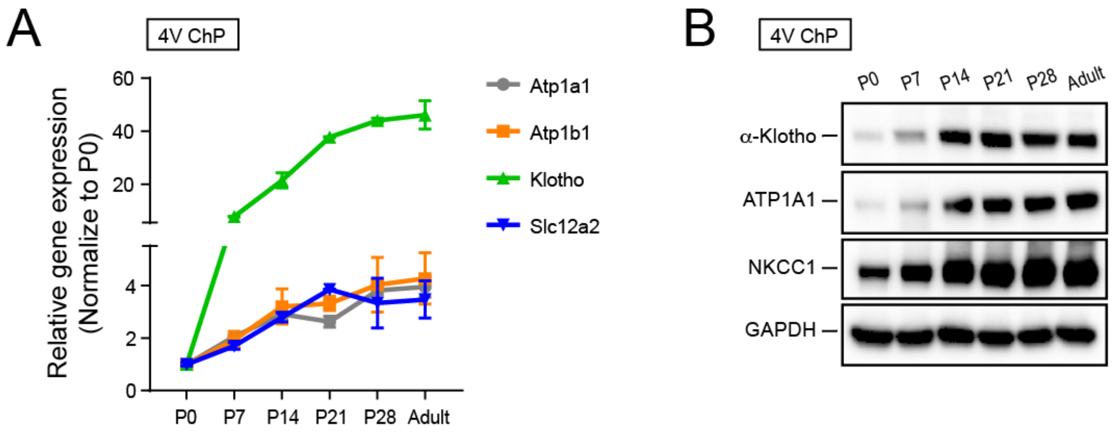


**Supplementary Fig 5. TRAP candidates validation in 4VChP. (A-B)** RT-qPCR analysis and immunoblotting analysis of 4V ChP during postnatal development.

*Supplementary figure 6*

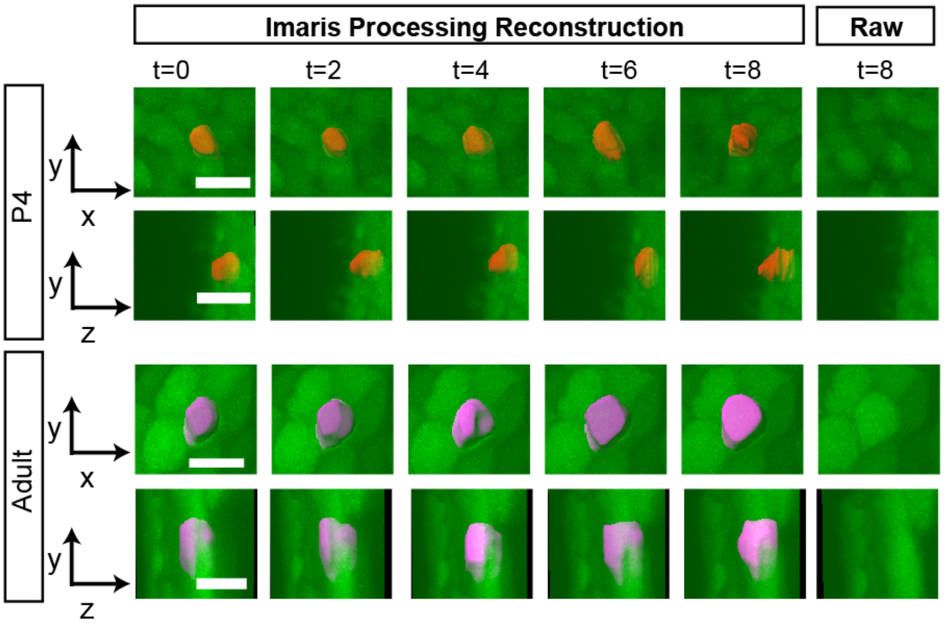


**Supplementary Fig 6.** Workflow of IMARIS demonstrating the cell volume quantification process. At each time point, the reconstructed 3D cell mask is highlighted (adult in pink and P4 in red) and views from x-y plane and y-z plane are displayed. Raw images from a single plane at the last time point are shown on the right end.

*Supplementary figure 7*

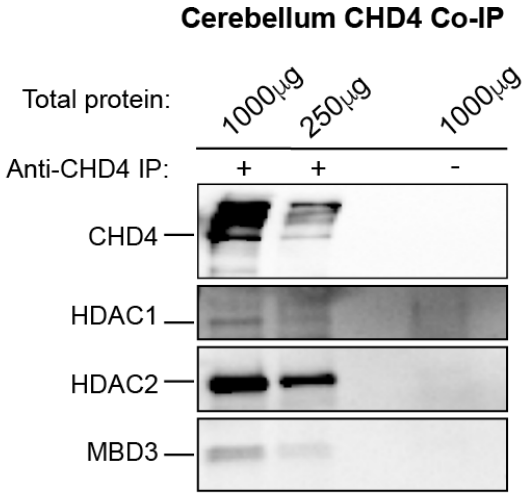


**Supplementary Fig 7.** Co-IP immunoblots using adult mouse cerebellum to validate the Co-IP protocol. An antibody targeting CHD4 co-immunoprecipitated several other complex members including HDAC1, HDAC2, and MBD3, from mouse cerebellar lysate, while a negative control performed with a control antibody of the same host species (in this case anti-NKCC1 antibody) failed to pull down any NuRD-CHD4 complex members.

*Supplementary figure 8*

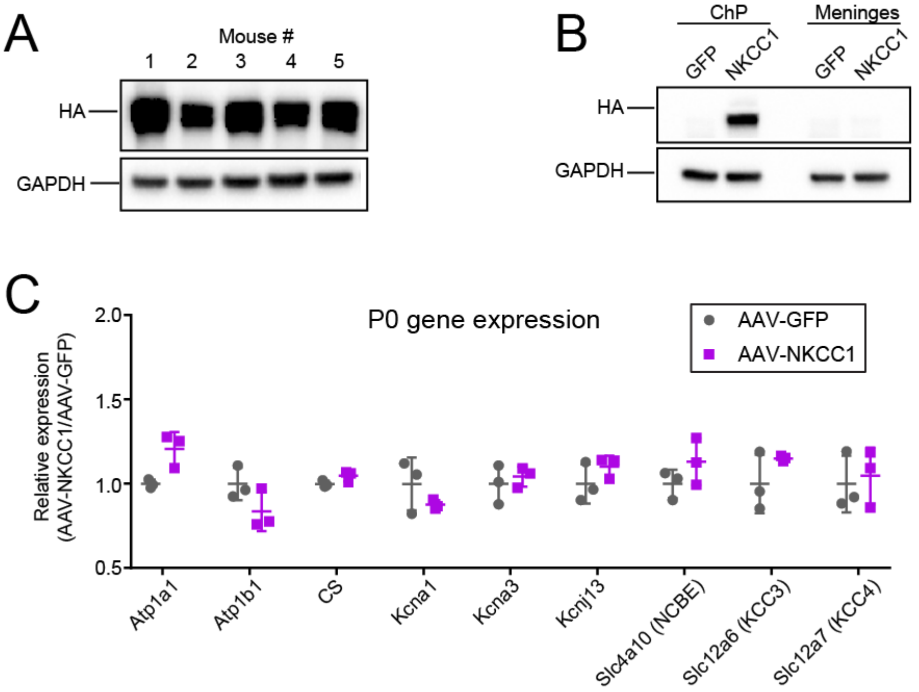


**Supplementary Fig 8. Validation of AAV2/5-NKCC1 transduction efficiency and specificity. (A)** Immunoblots of AAV2/5-NKCC1 transduced ChP showing successful but variable transduction rate within one litter. **(B)** Immunoblot of AAV transduced ChP and meninges showing non-detectable meningeal transduction by AAV2/5-NKCC1. **(C)** RT-qPCR analysis of all other $K^+$ transporters and channels in the ChP after in utero viral transduction, showing no significant changes; $\alpha > 0.05$, multiple t-test corrected for multiple comparisons using the Holm-Sidak method.

*Supplementary figure 9*

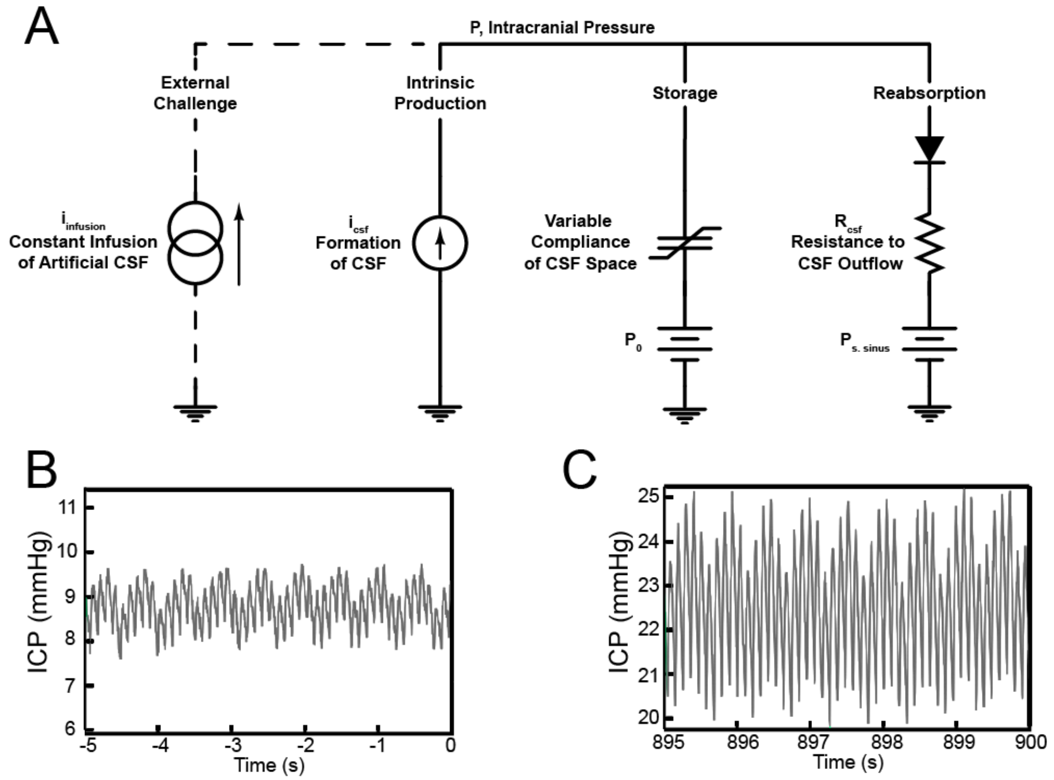


**Supplementary Fig. 9. Mechanisms of constant CSF infusion test by Marmarou's model.** **(A)** Marmarou's model of CSF dynamics. In this model, the physiologic processes of the cycle of CSF turnover are represented by analogous electric circuit elements, with ICP expressed as a solution to the circuit model in terms of lumped parameters describing the net effect of the processes on the level of ICP without attributing them to specific microscopic pathways. At the most basic level, the model is a statement of conservation of mass, with the rate of CSF production balanced by the rate of CSF storage in intracranial and spinal compartments plus the rate of CSF reabsorption. **(B-C)** Higher magnification of the ICP data of infusion test showing the normal arterial and respiratory components of the ICP waveform at the beginning of the test (**B**). The increase in waveform amplitude with ICP is expected with increasing volume load (**C**).

**Supplementary Table 1.** Summary of publications reporting various values of ChP epithelium intracellular $Na^+$, $K^+$, $Cl^-$ concentrations. (N.D., not determined)

| **Publication** | **$[Na^+]_i$** | **$[K^+]_i$** | **$[Cl^-]_i$** |
|---|---|---|---|
| Gregoriades, J. M. C., Madaris, A., Alvarez, F. J. & Alvarez-Leefmans, F. J. Genetic and pharmacological inactivation of apical Na(+)-K(+)-2Cl(-) cotransporter 1 in choroid plexus epithelial cells reveals the physiological function of the cotransporter. *American journal of physiology. Cell physiology* **316**, (2019) | *9.2± 2.5mM* | *ND* | *60.7± 12.3mM* |
| Steffensen, A. B. *et al.* Cotransporter-mediated water transport underlying cerebrospinal fluid formation. *Nat Commun* **9**, (2018). | *31 ± 5mM* | *141 ± 12mM* | *35 ± 9mM* |
| -Keep, R. F., Xiang, J. & Betz, A. L. Potassium cotransport at the rat choroid plexus. *The American journal of physiology* **267**, (1994).<br>-Zeuthen, T. The effects of chloride ions on electrodiffusion in the membrane of a leaky epithelium. Studies of intact tissue by microelectrodes. *Pflugers Archiv : European journal of physiology* **408**, (1987). | *30.0mM* | *120mM* | *50mM* |
| Johanson, C. E. & Murphy, V. A. Acetazolamide and insulin alter choroid plexus epithelial cell [Na+], pH, and volume. *The American journal of physiology* **258**, (1990). | *48 ± 0.7mM* | *95 ± 1.2mM* | *62±0.3mM* |
| Saito, Y. & Wright, E. M. Regulation of intracellular chloride in bullfrog choroid plexus. *Brain Res* **417**, (1987). | *10.5mM* | *ND* | *24mM* |
